# Maternal fluoxetine impairs synaptic transmission and plasticity in the medial prefrontal cortex and alters the structure and function of dorsal raphe nucleus neurons in offspring mice

**DOI:** 10.1101/2024.01.24.577094

**Authors:** Bartosz Bobula, Joanna Bąk, Agnieszka Kania, Marcin Siwiec, Michał Kiełbiński, Krzysztof Tokarski, Agnieszka Pałucha-Poniewiera, Grzegorz Hess

## Abstract

Selective serotonin (5-HT) reuptake inhibitors (SSRIs) are a class of antidepressant drugs commonly prescribed to women during pregnancy and breastfeeding to treat depression. There is evidence that prenatal exposure to SSRIs may be associated with a higher risk of adverse cognitive outcomes and affective disorders in later life. In animal models, exposure to SSRIs during brain development results in behavioral alterations as well as structural abnormalities of cerebral cortical neurons. Little is known about the consequences of SSRI-induced excess of 5-HT during development on the brain serotonergic system itself. In this study, an SSRI - fluoxetine (FLX) - was administered to C57BL/6J mouse dams during pregnancy and lactation. We found that maternal FLX decreased field potentials, impaired long-term potentiation, facilitated induction of long-term depression and tended to increase the density of 5-HTergic fibers in the medial prefrontal cortex (mPFC) of female but not male adolescent offspring. These effects were accompanied by deteriorated performance in the temporal order memory task and reduced sucrose preference with no change in marble burying behavior in FLX-exposed female offspring. We also found that maternal FLX reduced the axodendritic tree complexity of 5-HT dorsal raphe nucleus (DRN) neurons in female but not male offspring. Whole-cell recordings demonstrated no changes in the excitability of DRN 5-HT neurons in FLX-exposed offspring of either sex. While no effects of maternal FLX on inhibitory postsynaptic currents (sIPSCs) in DRN neurons were found, we observed a significant influence of FLX exposure on kinetic characteristics of spontaneous excitatory postsynaptic currents (sEPSCs) in DRN neurons. Finally, we report that no changes in field potentials and synaptic plasticity were evident in the mPFC of the offspring after maternal exposure during pregnancy and lactation to a new antidepressant, vortioxetine. These findings show that in contrast to the mPFC, long-term consequences of maternal FLX exposure on the structure and function of DRN 5-HT neurons are mild and suggest a sex-dependent, distinct sensitivity of cortical and brainstem neurons to FLX exposure in early life. Regarding side effects on brain development, vortioxetine might be a safer alternative to FLX.

## 1. Introduction

Between 10 – 20% of pregnant women suffer from depression. During and following pregnancy most of diagnosed women use antidepressant drugs (reviewed in: Hayes et al., 2012; Lebin and Novick, 2022), mainly the selective serotonin (5-hydroxytryptamine, 5-HT) reuptake inhibitors (SSRIs) that inhibit presynaptic 5-HT transporter (Hiemke and Härtter, 2000). There is controversy whether exposure to SSRIs early in life is safe or it affects the health of the child. Recent reports show that antidepressant drug use during pregnancy does not increase the risk of major negative outcomes, including neurodevelopmental disorders, in progeny (reviewed in: Besag and Vasey, 2023; Suarez et al., 2022). However, a substantial body of evidence indicates that prenatal exposure to SSRIs may lead to adverse cognitive outcomes and affective disorders (reviewed in: Desaunay et al., 2023; Lebin and Novick, 2022). In particular, prenatal exposure to SSRIs has been associated with a higher risk of autism spectrum disorder in later life (Andalib et al., 2017; Schendel, 2017). One of the most commonly prescribed SSRIs is fluoxetine (FLX; Kiryanova et al., 2013; Molenaar et al., 2020). It has been shown that FLX can transfer to the fetus through the placenta, and has also been found in breast milk (Heikkinen et al., 2003, Kim et al., 2006). During brain development 5-HT acts as a main modulator of neuronal proliferation, migration, differentiation, synaptic plasticity, and apoptotic cell death (reviewed in: Higa et al., 2022; Teissier et al., 2015; Vitalis and Parnavelas, 2003). Thus, FLX-induced increase in tissue 5- HT content may interfere with the normal development of brain circuitry (reviewed in: Adjimann et al., 2021).

Experiments on rodents have confirmed that FLX can transfer from the organism of the dam to the offspring (Kiryanova et al., 2013; Maloney et al., 2018). A number of animal studies investigated the consequences of maternal FLX exposure on offspring behavior in later life (reviewed in: Glover and Clinton, 2016; Ramsteijn et al., 2020), however only a handful of them examined the consequences for the offspring when dams were exposed to FLX during the whole pregnancy and lactation period. FLX exposure is often related to decreased exploratory locomotion in the offspring as well as deficits in social and sexual behaviors, and occurrence of anxiety-and depression-like phenotypes (reviewed in: Kiryanova et al., 2013; Kinast et al., 2013). Other studies reported decreased anxiety-like and depression-like behavior (McAllister et al, 2012), increased aggressive behavior (Kiryanova et al., 2016), social communication and interaction deficits as well as occurrence of repetitive patterns of behavior (Maloney et al., 2018). There are sex differences in the effects of perinatal FLX exposure on the behavior of the offspring in adolescence (Lisboa et al., 2007; Ramsteijn et al., 2020). These behavioral alterations have been linked to disturbances in glutamatergic transmission in the prefrontal cortex and hippocampus, including decreased expression of NMDA receptor, metabotropic glutamate receptor 1 (mGluR1) and postsynaptic density protein 95 (PSD-95) that coincide with the anxiety-like and depression-like phenotype (Millard et al, 2019). It has recently been reported that maternal FLX exposure results in a decrease in the dendritic complexity and spine density of pyramidal neurons in the medial prefrontal cortex (mPFC) of the offspring mice (Maloney et al., 2022). However, it is not known if and how maternal exposure to FLX throughout gestation and lactation affects synaptic transmission and plasticity in the offspring mPFC.

In the adult organism the 5-HT system is involved in multiple cognitive and physiological processes including sleep, cognition, sensory perception, motor activity, temperature regulation, appetite, hormone secretion, nociception and sexual behavior (reviewed in: Jacobs and Azmitia, 1992). 5-HT also modulates synaptic plasticity in the prefrontal cortex (Zhong et al., 2008a; reviewed in: Ruggiero et al., 2021). Widespread, long-range 5-HT projections to the forebrain originate from the dorsal raphe nucleus (DRN; termed also the B7 nucleus: Muzerelle et al, 2016; Sargin et al, 2019; Silva et al., 2010). 5-HT neurons projecting to the mPFC are located in the rostral and central subregions of the DRN (Chandler et al., 2013). Given the role that the 5-HT system plays in brain functioning, it is important to understand the consequences of SSRIs exposure in early life on the 5-HT system itself. However, the influence of maternal exposure to SSRIs throughout the whole gestation and lactation period on the electrophysiology and morphology of the offspring DRN neurons has not yet been investigated.

Thus, in the present study we first verified the consequences of maternal exposure to FLX delivered in drinking water throughout gestation and lactation on synaptic transmission and induction of long-term synaptic plasticity in *ex vivo* slices of the offspring mPFC using field potential recordings. Next, using a combination of whole-cell recordings from *ex vivo* brainstem slices and morphological analyses of recorded cells we investigated the effects of exposure of dams to FLX on the electrophysiological and morphological characteristics of identified DRN 5-HT neurons of the male and female offspring.

In contrast to FLX, a newer generation antidepressant - vortioxetine (VOR) - not only inhibits the 5-HT transporter but also acts as an agonist of the 5-HT1A receptor, partial agonist of the 5-HT1B receptor, and an antagonist of 5-HT1D, 5-HT3 and 5-HT7 receptors (Sanchez et al., 2015). VOR is considered an effective switch therapy for patients whose response to SSRIs is inadequate (Thase et al., 2017). In the final step of this study we compared the effects of maternal exposure to VOR on synaptic transmission and long-term potentiation in offspring mPFC slices with those following the exposure to FLX.

## 2. Material and methods

### 2.1. Animals and treatment

Experiments were carried out in accordance with the European Communities Council Directive of September 22, 2010 (2010/63/UE) on the protection of animals used for scientific purposes and national law. Experiments were approved by the 2^nd^ Local Institutional Animal Care and Use Committee at the Maj Institute of Pharmacology, Polish Academy of Sciences in Krakow. C57BL/6J mice (Charles River Laboratories, Sulzfeld, Germany) were housed on a 12/12 h light/dark cycle with light on between 7:00 and 19:00 hours and with standard food (RM3, Special Diet Services, Witham, UK) available *ad libitum*. Mice had free access to tap water before mating and the average daily drinking volume was recorded. Mice were mated at the age of 75 – 90 days (1 female and 1 male per cage) at the end of the day and females were checked the following day for the presence of a vaginal plug (the gestational day 0, GD 0). Pregnant mice were housed singly.

During pregnancy and lactation dams of the experimental group received FLX (TCI Europe NV, Zwijndrecht, Belgium) with drinking water (concentration: 0.045 mg/ml) at a daily dose of approx. 7.5 mg/kg based on the weight of animals on GD 0 (Lisboa et al., 2007; Navailles et al., 2008; Oh et al., 2009). Dams began to receive FLX in drinking water only after confirming the pregnancy since it has been shown that FLX may dysregulate estrous cycle in mice (Domingues et al., 2023). In a subset of experiments dams received VOR (purchased from Lundbeck, Valby, Denmark) that was incorporated into rodent chow at a concentration of 1.8 g/kg of food weight (RM3, Special Diet Services, Witham, UK). This amount corresponds to a daily dose of VOR of approximately 10 mg/kg (Felice et al., 2018). FLX or VOR delivery was discontinued on postnatal day (PND) 18. Dams of the control group had free access to tap water and food. Offspring of experimental and control dams were weaned on PND 25 and group-housed by sex with free access to standard food and tap water.

### 2.2. Behavioral tests

#### Temporal order memory task

**(TOMT)** The experiment was carried out using black boxes of 42 x 30 x 22 cm, covered with sawdust and illuminated by a 25 W bulb, placed in a dark room. The task consisted of two habituation trials, performed on two consecutive days, and the test trial, performed on the 3^rd^ day (Pałucha-Poniewiera et al., 2023). During habituation trials each mouse was placed individually in the box without objects and allowed to explore the environment for 10 min. Twenty-four hours later, the proper test comprising two training phases (TP1 and TP2) with an interval of 3 hours and one test trial were performed. In each training phase mice were allowed to explore two copies of an identical object for 4 min. During TP1, two X objects were used, and during TP2, two Y objects were used. One hour after TP2, the test trial lasting 3 min was performed. During this phase, a third copy of object X and a third copy of object Y were used. If temporal order memory is intact mice spend more time exploring (i.e., sniffing or touching) object X than object Y. Time (T) spent exploring objects X and Y was measured by an experienced observer, followed by calculation of the discrimination index [(TX – TY)/(TX + TY)].

#### Sucrose preference test

**(SPT)** The sucrose preference test was performed according to Pałucha-Poniewiera et al (2021) with some modifications. After being individually caged, mice were given a choice between a bottle with 1% sucrose solution and another bottle with tap water for 48 hours. The position of the bottles was switched every 12 hours. At the beginning and the end of the test the bottles were weighed, and liquid consumption was calculated. The preference for sucrose was calculated as a percentage of consumed sucrose solution in terms of the total amount of the liquid drunk.

#### Marble burying test

**(MBT)** Polycarbonate cages (26 x 48 x 20 cm) were filled with a 4.5 cm layer of standard bedding, followed by gently overlaying 20 glass marbles (15 mm diameter) in a regular 4 × 5 arrangement (Deacon, 2006). Mice were placed individually inside cages for a 30-minute exploration period. Then, the number of marbles buried (defined as more than 50% of the marble covered by bedding) was recorded. White noise (55 dB) was present during testing to dampen ambient noise.

### 2.3. Field potential recording and LTP induction in slices of the medial prefrontal cortex

Brain slices of the frontal cortex (380 μm thick) were prepared from FLX- or VOR- exposed and control male and female offspring aged 8 – 12 weeks (PND 56 – 80). Coronal slices were cut using a vibrating microtome (Leica VT1000s) in ice-cold artificial cerebrospinal fluid (NMDG-HEPES ACSF) containing (in mM): NMDG (92), KCl (2.5), CaCl2 (0.5), MgSO4 (10), NaH2PO4 (1.2), NaHCO3 (30), HEPES (20), glucose (25), sodium ascorbate (5), thiourea (2), sodium pyruvate (3), pH titrated to 7.3 with HCl. Slices were then incubated at 35°C for 25 min in NMDG-HEPES aCSF while gradually introducing NaCl (the ‘Na spike-in’ method (Ting et al., 2018). Afterwards, slices were transferred to room-temperature recording ACSF (see below).

Individual slices were placed in the recording chamber of the interface type and were superfused with modified ACSF containing (in mM): NaCl (132), NaHCO3 (26), KCl (2), CaCl2 (2.5), MgSO4 (1.3), KH2PO4 (1.25) and D-glucose (10), bubbled with 95% O2 – 5% CO2 at 2.5 ml/min (32 ± 0.5 °C). Field potentials (FPs) were recorded using a glass micropipette filled with ACSF (1–2 MΩ), that was placed in cortical layer II/III of the prelimbic (PL) subdivision of the mPFC. FPs were evoked with stimuli (duration: 0.2 ms) applied at 0.033 Hz via a constant-current stimulus isolation unit (WPI) using a concentric Pt- Ir stimulating electrode (FHC) that was placed in underlying sites in layer V. FPs were amplified (Axoprobe 1A, Axon Instruments), A/D converted at 10 kHz and stored using Micro1401 interface and Signal 2 software (CED).

A stimulus–response curve was made for each slice. To obtain the curve, stimulation intensity was gradually increased stepwise (5–100 μA) and one response was recorded at each intensity. The stimulus–response curves obtained for each slice were fit with the Boltzmann equation: Vi = Vmax/(1 + exp((u − uh)/−S), where Vmax is the maximum FP amplitude; u is the stimulation intensity; uh is the stimulation intensity evoking FP of half-maximum amplitude; S is the factor proportional to the slope of the curve. For the induction of LTP theta burst stimulation (TBS) protocol was used (Popek et al., 2020). Before TBS, stimulation intensity was adjusted to evoke a response of 30% of the maximum FP amplitude. Standard TBS protocol consisted of 10 trains of stimuli at 5 Hz, each composed of five pulses at 100 Hz, repeated five times every 15 s. The amount of LTP was determined as an average increase in the amplitude of FPs recorded between 45 and 60 min after TBS, relative to baseline. In some experiments a “weak” TBS protocol was used, during which 10 trains of stimuli were repeated two instead of five times.

### 2.4. Whole-cell recording from brainstem slices

Brainstem coronal slices containing a part of the DRN (250 μm thick) were prepared from FLX-exposed and control male and female offspring aged 8 – 12 weeks (PND 56 – 80), similarly to frontal cortex slices (see above). An individual slice was placed in the recording chamber and perfused at 2.5 ml/min with warm (32 ± 0.5°C) ACSF-containing (in mM): NaCl (132), NaHCO3 (26), KCl (2), CaCl2 (2.5), MgSO4 (1.3), KH2PO4 (1.25) and D-glucose (10), bubbled with 95% O2 – 5% CO2. Patch micropipettes (open tip resistance of approx. 6 MΩ) were pulled from borosilicate glass capillaries (Harvard Apparatus), using the Sutter Instruments P97 puller and filled with the solution containing (in mM): K-gluconate (130), NaCl (5), CaCl2 (0.3), MgCl2 (2), HEPES (10), phosphocreatine-Na2 (10), Na2-ATP (5), Na-GTP (0.4) and EGTA (0.3). Osmolarity of the solution was 290 mOsm, and pH was adjusted to 7.2 with KOH. Cells in the DRN midline region were visualized using the Zeiss Axioskop 1 microscope with Nomarski optics, a 40× water immersion lens and an infrared camera.

Signals were acquired with SEC 05LX amplifier (NPI), low-pass filtered at 2 kHz and digitized at 20 kHz using the Digidata 1440A interface and pClamp 10 software (Molecular Devices). Putative 5-HT neurons were identified on the basis of their characteristic response to a series of hyper-and depolarizing current pulses (duration: 500 ms) in 20 pA increments (from ™40 to 140 pA; Sowa et al., 2018). The relationship between the injected current intensity and number of evoked action potentials was plotted for each cell and the gain was defined as a slope of the linear regression line fitted to experimental data. Total (cumulative) count of spikes evoked by all ten current pulses was also calculated for each neuron.

Next, cells were clamped at either −76 or 0 mV for recording of sEPSCs or sIPSCs, respectively. After 15 min of stabilization synaptic events were recorded for 4 min. Individual synaptic events were detected offline using the Mini Analysis program (Synaptosoft) and manually selected for further analysis. Only recordings during which the access resistance ranged between 15 – 18 MΩ and remained stable throughout the recording (< 25% change) were accepted for analysis. The thresholds for the detection of EPSCs and IPSCs were 6 pA and 10 pA respectively. The kinetic parameters characterizing synaptic currents were determined from averaged waveforms. The rise time was defined at the time needed for the current to rise from 10% to 90% of the peak. The decay time constant (tau) was determined by fitting an exponential function to the decay phase of the waveform.

In a subset of slices the effect of the activation of 5-HT1A receptors in DRN neurons was tested in the voltage-clamp mode at a holding potential of ™60 mV. The magnitude of the outward current evoked by bath application of 5-carboxamidotryptamine (5-CT, 1 μM) lasting 1 min was measured (Beck et al., 2004).

### 2.5. Post-recording immunostaining of brainstem slices

To visualize cells filled with biocytin, after successful recordings the slices were initially fixed in a 4% solution of paraformaldehyde in PBS at RT for one hour. Following several rounds of washing, the slices were incubated overnight at RT in a PBS solution containing 0.1% Triton X100, and specific goat antibodies targeting TPH-2 (1:1000, ab121013, Abcam, UK). Afterwards slices were repeatedly washed and incubated overnight at RT in PBS containing a mixture of Alexa Fluor 647-conjugated streptavidin (1:500, S21374, Invitrogen, UK) and secondary antibodies (donkey anti-goat Alexa Fluor Plus 488, 1:500, A32814, Invitrogen, UK). Following another round of washing, the slices were mounted onto glass slides, sealed with Vectashield mounting medium (Vector Laboratories, Newark, USA) and coverslipped. Optical z-stacks were then captured with a 20x/0.8 air objective and a 63x/1.4 oil-immersion plan apochromatic objective on a Zeiss Axio Imager Z2 fluorescence microscope equipped with the Apotome 2 optical sectioning module (Carl Zeiss, Germany), to gather data on individual neuron morphology and immunohistochemistry.

### 2.6. Assessment of dendritic morphology

TPH2-immunoreactive (TPH2-ir) neurons filled with biocytin during recording with complete, well-stained and clearly visible dendritic trees were imaged using Zeiss AxioImager Z2 fluorescent microscope equipped with the Apotome 2 optical sectioning module, Axiocam 506 camera and Colibri 5/7 light source (Carl Zeiss, Germany), under 20x/0.8 air objective. Nyquist-bounded z-stacks of images were obtained for the entire visible dendritic arbor of the imaged neuron. Subsequent tracing of neuronal branches combined with 3D Sholl analysis (10 μm step size) was performed on the acquired images in ImageJ (Fiji, RRID:SCR_002285; Schneider et al., 2012) with the Simple Neurite Tracer plugin (RRID:SCR_016566; Longair et al., 2011). Other parameters describing the axodendritic tree of cells (branch length, number of primary branches, total number of branches, number of branch points, number of branch tips and maximum branch order) were obtained with the same software.

### 2.7. Tissue preparation and immunostaining for 5-HT

Mice were transcardially perfused with 15 ml of ice-cold PBS, followed by 30 ml of cold PBS with 4% paraformaldehyde (PFA), prepared fresh from 16% methanol-free stock (28906, Thermo Fisher Scientific, Poland). Brains were extracted and postfixed overnight in PBS with 4% PFA at 4°C. The next day, 50 µm-thick coronal slices containing the mPFC were cut on a vibrating microtome (vt1200S, Leica Microsystems, Germany). Following 3×10 min washes in PBS at RT, the slices were incubated for 48h at 4°C in PBS containing 0.1% Triton X-100 and a 1:1000 dilution of a goat primary antibody against serotonin (20079, Immunostar, USA). After 3×10 min washes in PBS slices were incubated for 2h at RT in PBS containing a secondary antibody (donkey anti-goat Alexa Fluor Plus 647, 1:500, A21447, Invitrogen, UK). During a final round of washes tissue slices were counterstained with DAPI (NucBlue™ Fixed Cell ReadyProbes™ Reagent, R37606, Invitrogen, UK) according to the kit manufacturer’s instructions. Slices were then mounted on glass slides in Prolong Glass medium (P36980, Invitrogen, UK), coverslipped and cured.

### 2.8. Image acquisition and analysis

Slides (n = 3 animals per group; 3-5 replicate sections per animal) were imaged on Zeiss AxioImager Z2 fluorescent microscope equipped with an ApoTome module, Axiocam 506 detector and Colibri 5/7 light source, under 63x oil objective (Plan Apochromat 1.4 M27, Zeiss). Photomicrographs were taken bilaterally from three anatomical areas: motor cortex (corresponding to M2, dorsal part), prelimbic (PL) cortex and infralimbic (IL) cortex. At each position, stacks of images (optical slice z = 0.32 µm) spanning the innermost 8 µm of the tissue were obtained, with 10 Apotome slices per frame (i.e. a total of 260 images per site). Illumination intensity and exposure times were adjusted automatically for each image series to obtain 80% coverage of the camera’s dynamic range.

Following acquisition, the image series were deconvolved using proprietary Zeiss Zen 2.6 software (ApoTome deconvolution, automatic normalization with background and fluorescent decay corrections). Subsequently, segmentation and analysis were performed automatically in ImageJ software (Schindelin et al., 2012). Images were filtered by background subtraction (rolling ball radius = 5 pixels) followed by smoothing; maximum intensity projections were then created from the resulting filtered image stacks. These were then automatically locally segmented with the Phansalkar method (with the following parameters, empirically adjusted on example images from the dataset: r = 15, a = 0.1, b = 0; Phansalkar et al., 2011) and the obtained segmented objects were high-pass-filtered to remove debris and any residual noise (size > 15 pixels). Within each cortical area, outer regions, corresponding to cortical layers I-III and inner regions (layers V-VI) were manually delineated and analyzed separately. The resulting measurement was the area of the segmented 5-HT-immunopositive processes in each region of interest, expressed as a percentage of the total area of the region (Area Fraction, AF).

### 2.9. Statistical analysis

Statistical analysis of electrophysiological data was carried out using two-way analysis of variance (ANOVA) with FLX exposure and sex considered as factors. Tukey’s or Sidak’s post hoc tests were used if the interaction was statistically significant (*p* < 0.05).

The results of behavioral tests were analyzed using unpaired, two-tailed Student’s t-test. Differences between groups were considered significant if the *p*-values were < 0.05.

For the analysis of 5-HT-immunopositive fibers AF measurements corresponding to each region of interest, i.e. combination of anatomical structure (prelimbic, infralimbic, or motor cortex) and depth (superficial and deep layers) were averaged between sides (left-right) and then between technical replicates taken from each animal. A linear model was then fitted using R *lm* function (R Core Team, 2022), with average AF as the dependent variable and group (control females, FLX females, control males, FLX males) as well as location (all combinations of anatomical site and depth) as the independent variables. For the group variable, a contrast matrix of the form:

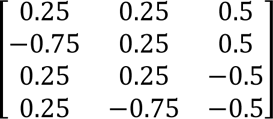

was used, resulting in the following planned comparisons:

- [,1] con F vs. FLX F; estimating the difference in AF between control females and FLX- exposed females
- [,2] con M vs. FLX M; estimating the difference in AF between control males and FLX-exposed males
- [,3] con F vs. con M; estimating baseline differences in AF between control female and male mice

For the location variable, deviation (sum) contrasts were used. Model viability was initially assessed by visual inspection of diagnostic plots (not shown). The AIC metric was then used for comparing fitted models and ANOVA was used for testing the significance of specific terms, confidence intervals and corresponding *p*-values for marginal effects were used for group comparisons. Data were considered to be significantly different at *p* < 0.05.

All analyses were *performed* using the GraphPad Prism for Windows software version 10.0.2 (GraphPad Software) unless indicated otherwise.

## 3. Results

### 3.1. Maternal fluoxetine decreases field potentials and impairs long-term potentiation in the mPFC of female but not male offspring

To verify the effects of the procedure of maternal exposure to FLX we began with extracellular recordings of FPs from layer II/III of the PL subdivision of mPFC in *ex vivo* slices of the offspring brain. There was a significant main effect of FLX exposure (two-way ANOVA, F(1, 45) = 10.36, *p* = 0.0024) and a significant main effect of sex (F(1, 45) = 11.56, *p* = 0.0014) on the maximum amplitude of FPs calculated using Boltzmann fits (Fig. 1A, B), but no significant interaction between treatment and sex factors (F(1, 45) = 3.884, *p* = 0.0549). Mean maximum amplitudes of FPs (± SEM) in 4 tested groups were as follows: FLX-exposed females (termed: FLX F): 0.71 ± 0.04 mV, n = 12; control females (termed: con F): 0.99 ± 0.06 mV, n = 12; FLX-exposed males (termed: FLX M): 1.0 ± 0.07 mV, n = 12; control males (termed: con M): 1.07 ± 0.05 mV, n = 13 (Fig. 1C).

**Fig. 1.**
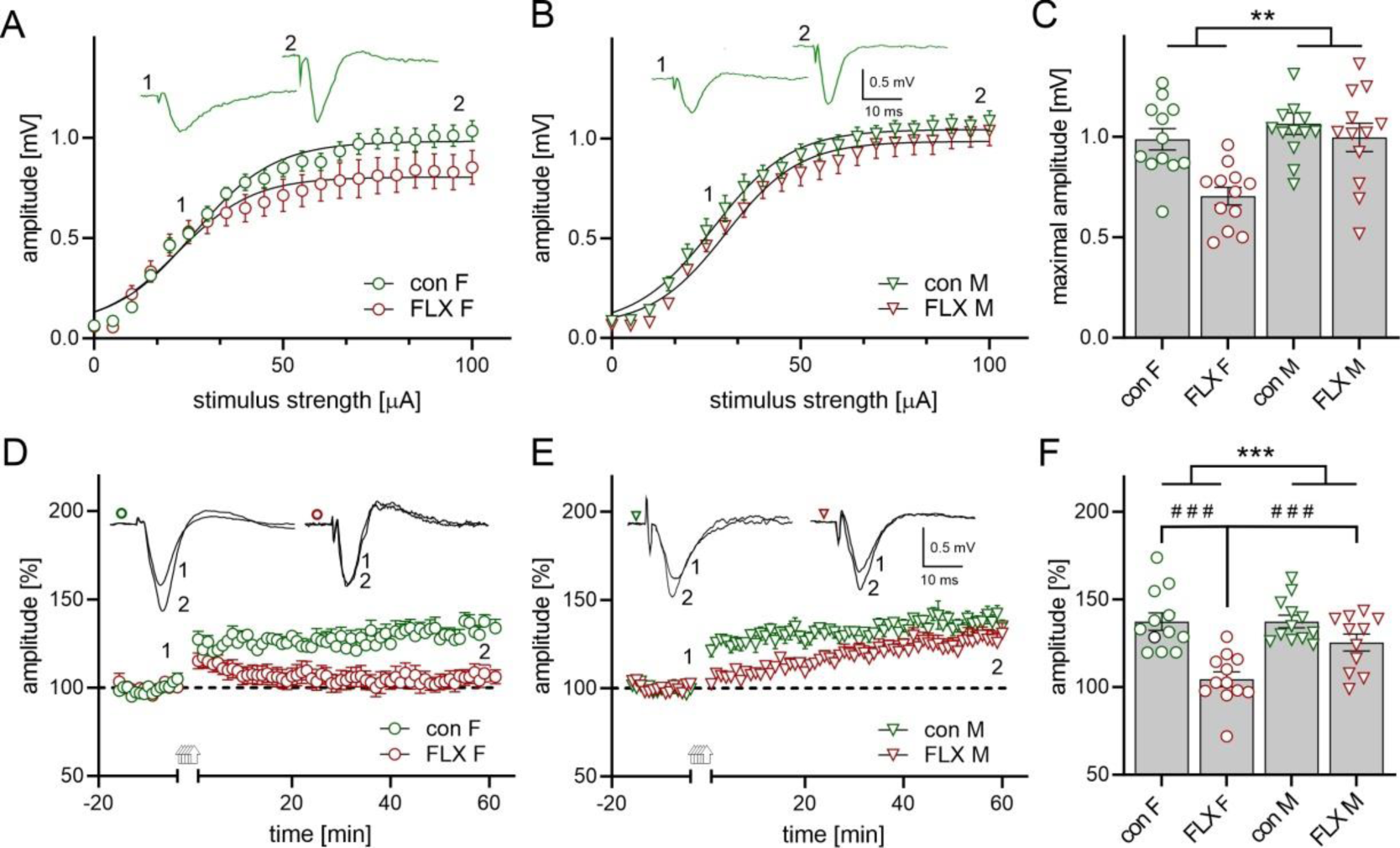
Maternal FLX reduces the amplitude of FPs and impairs LTP in the mPFC of female but not male offspring. Graphs (A, B) illustrate the effect of FLX exposure on the relationship between stimulus intensity and mean FP amplitude (± SEM) in mPFC slices obtained from female (F, circles) and male (M, triangles) offspring that were exposed to FLX (red symbols) and control mice (con, green symbols). Insets in A and B show representative FPs from control mice at two stimulation intensities (1, 2). Black lines represent fits to the Boltzmann equation (see: Material and methods). (C) Comparison of calculated maximum FP amplitudes in four groups of animals. Shown are mean values ± SEM with circles and triangles representing individual slices. Two-way ANOVA, main effect of FLX exposure: *p* < 0.01, main effect of sex: \*\**p* < 0.01, FLX exposure x sex interaction: *p* > 0.05. Symbols and labels as in A, B. (D, E) Plots of the amplitude of FPs (mean ± SEM) recorded from mPFC slices obtained from control (con F, green circles) and FLX-exposed females (FLX F, red circles) as well as from control (con M, green triangles) and FLX-exposed males (FLX M, red triangles). Arrows denote the time of theta-burst stimulation (TBS, repeated 5 times). Insets in D and E show superposition of FPs recorded in representative experiments before and after TBS at times indicated by numbers. (F) Mean (± SEM) amplitude of FPs recorded between 45 – 60 min after TBS, relative to baseline. Two-way ANOVA, main effect of FLX exposure: *p* < 0.05, main effect of sex: \*\*\**p* < 0.0001, FLX exposure x sex interaction: *p* < 0.005, multiple comparison test: ###*p* < 0.0001.

Two-way ANOVA revealed a significant main effect of FLX exposure (F(1, 43) = 5.498, *p* = 0.0237), a significant main effect of sex (F(1, 43) = 27.27, *p* < 0.0001), and an interaction between FLX exposure and sex (F(1, 43) = 5.821, *p* = 0.0202) on the amplitude of FPs recorded 45 - 60 min after stimulation of the slice with the standard TBS protocol relative to baseline. Induction of LTP was impaired in slices obtained from FLX-exposed female offspring compared to the control group (Fig. 1D; FLX F vs. con F: 104.3 ± 4.2 %, n = 12 vs. 137.3 ± 5.0 %, n = 12, respectively; q = 7.721, *p* < 0.0001; Tukey’s multiple comparisons test). In contrast, although the time course of the change in FP amplitude over 1 hour after stimulation with 5 trains of TBS in slices obtained from FLX-exposed male offspring and control animals was different (Fig. 1E), at the end of the recording the LTP magnitude did not differ between slices obtained from FLX-exposed and control mice (FLX M vs. con M: 125.4 ± 4.8%, n = 11 vs. 137.4 ± 3.3 mV, n = 12, respectively; q = 2.778, *p* = 0.2172; Tukey’s test; Fig. 1F). We conclude that maternal FLX exposure resulted in a selective LTP impairment in female offspring (FLX M vs. FLX M: q = 4.705, *p* = 0.0094; Tukey’s test; Fig. 1F).

### 3.2. Maternal fluoxetine alters behavior of female offspring

To explore the potential behavioral correlates of the observed effect of maternal exposure to FLX on LTP in mPFC of female offspring a series of behavioral tests was conducted in FLX-exposed and control females. Analysis of the performance of mice in the TOMT test showed a significant reduction of the discrimination index in the FLX-exposed group compared to the control group (FLX F vs. FLX con: t = 7.081, df = 22, *p* < 0.0001; t-test; Fig. 2A1), which suggests a disturbance of the temporal order memory induced by FLX exposure.

**Fig. 2.**
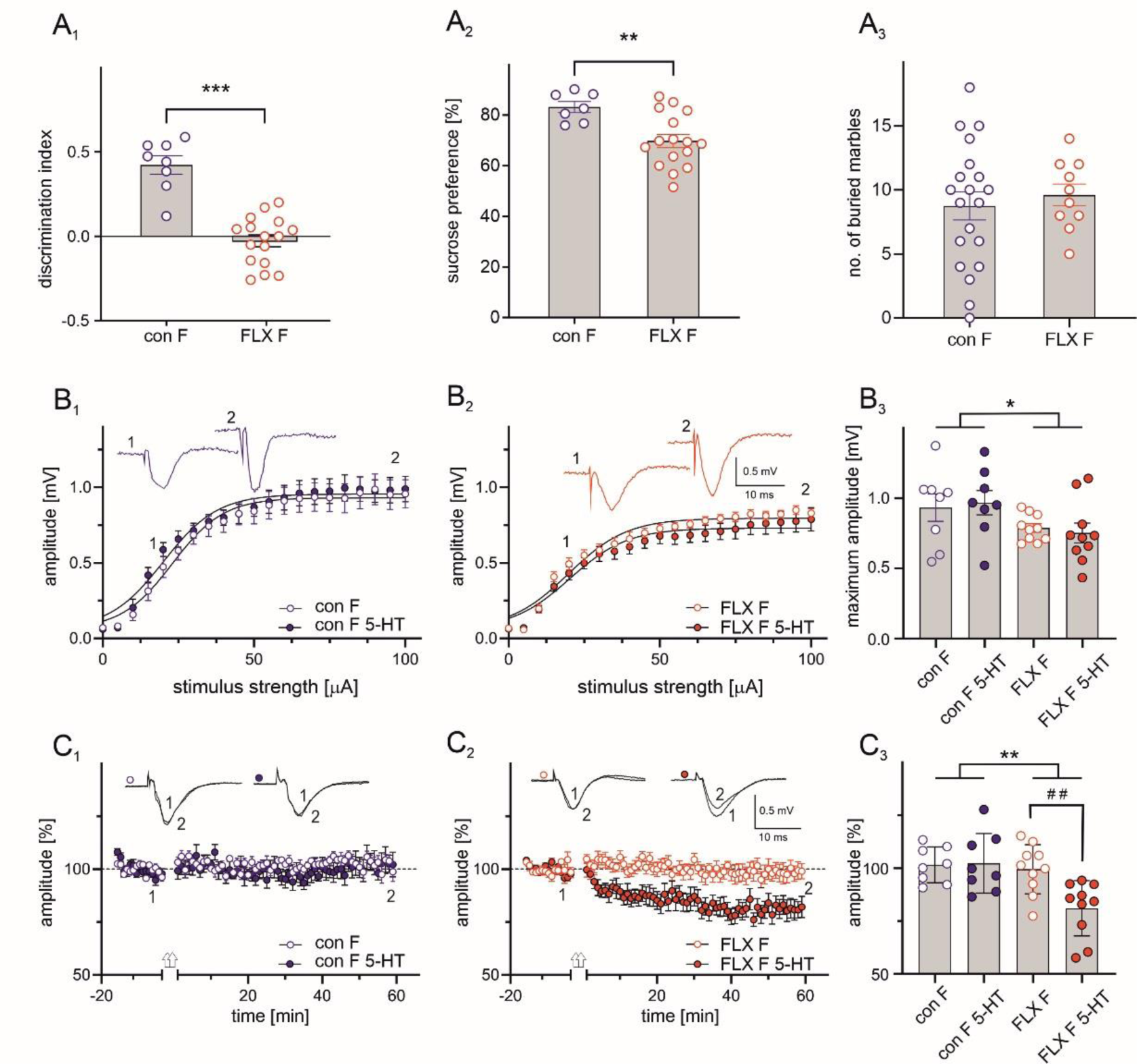
Maternal FLX induces behavioral deficits and modifies the 5-HT-dependent threshold for long-term depression in the mPFC of female offspring. Effects of maternal FLX on the discrimination index in the temporal order memory task (A1), sucrose preference (A2), and marble burying test (A3). Shown are mean values ± SEM with circles representing individual animals; con F – control; FLX F – fluoxetine-exposed female mice; *** *p* < 0.001; \*\**p* < 0.01, t-test. (B1) The relationship between stimulus intensity and mean FP amplitude (± SEM) in slices obtained from control female mice (con F) recorded in standard ACSF (open circles) and in ACSF supplemented with 5 µM 5-HT (5-HT; filled circles). Insets show examples of representative FPs at two stimulation intensities (1, 2). Lines represent fits to the Boltzmann equation. (B2) Same as in B1 but FPs were recorded from slices obtained from FLX-exposed females. (B3) Comparison of maximum calculated FP amplitudes in four groups of animals. Shown are mean values ± SEM with circles representing individual slices. Two-way ANOVA, main effect of FLX exposure: \**p* < 0.05, main effect of 5-HT: *p* > 0.05, FLX exposure x 5-HT interaction: *p* > 0.05. (C1, 2) plots of the mean amplitude of FPs (± SEM) recorded before and after “weak” TBS (repeated 2 times) from slices obtained from control (C1) and FLX-exposed female mice (C2) in standard ACSF (open circles) and in the presence of 5 µM 5-HT (filled circles). Arrows denote the time of “weak” TBS (repeated 2 times). Insets in C1 and C2 show superposition of FPs recorded in the course of representative experiments before and after “weak” TBS at times indicated by numbers. (C3) mean (± SEM) amplitude of FPs recorded between 45-60 min after “weak” TBS, relative to baseline. Two-way ANOVA, main effect of FLX exposure: \*\**p* < 0.01, main effect of 5-HT present in ACSF: *p* < 0.05, FLX exposure x 5-HT in ACSF interaction: *p* < 0.05, multiple comparisons: ##*p* < 0.01.

Decreased sucrose preference was evident in FLX-exposed mice compared to the control group (FLX F vs. FLX con: t = 3.117, df = 21, *p* = 0.0052; t-test; Fig. 2A2) in the SPT test, suggestive of an anhedonia-like state in these animals.

In contrast, no difference in the number of buried marbles between FLX-exposed and control mice was observed in the MBT test (FLX F vs. FLX con: t = 0.5112, df = 28, *p* = 0.6132; t-test; Fig. 2A3).

### 3.3. Maternal fluoxetine and exogeneous 5-HT modify the threshold for long-term depression in the mPFC of female offspring

Since maternal exposure to FLX impaired LTP in the PL subdivision of the mPFC in female offspring, we next tested whether maternal exposure to FLX can also affect long-term depression (LTD). In a subset of mPFC slices obtained from female offspring a “weak”, subthreshold TBS protocol involving 2 repetitions of TBS (instead of 5), was employed. Since exogenous 5-HT may facilitate induction of LTD in mPFC preparations (Zhong et al., 2008a), we incubated those slices either in standard ACSF or in ACSF supplemented with 5- HT in low concentration (5 µM). Comparison of Boltzmann fits confirmed that there was a significant main effect of FLX exposure (two-way ANOVA, F(1, 35) = 6.59, *p* = 0.0151) on the maximum amplitude of FPs (Fig. 2B1 - B3) while there was no significant main effect of the addition of 5-HT to the ACSF on FPs (F(1, 32) = 0.0005, *p* = 0.9832) and no interaction between FLX exposure and the presence of 5-HT in ACSF (F(1, 32) = 0.1625, *p* = 0.6896). Mean maximum amplitudes of FPs (± SEM) recorded in standard ACSF were as follows: control females (con F): 0.93 ± 0.09 mV, n = 8; FLX-exposed females (FLX F): 0.78 ± 0.03 mV, n = 10. Mean maximum amplitudes of FPs (± SEM) recorded in ACSF supplemented with 5-HT were as follows: control females (con F 5-HT): 0.96 ± 0.08 mV, n = 8; FLX-exposed females (FLX F 5-HT): 0.75 ± 0.07 mV, n = 10.

Two-way ANOVA revealed a significant effect of FLX exposure (F(1, 32) = 5.498, *p* = 0.0069), a significant effect of the addition of 5 µM 5-HT to ACSF (F(1, 32) = 4.836, *p* = 0.0352), and a significant interaction between FLX exposure and supplementation of ACSF with 5-HT (F(1, 32) = 5.628, *p* = 0.0239) on the amplitude of FPs recorded between 45 - 60 min after stimulation with the “weak” TBS (2 trains), relative to baseline. In slices obtained from control female offspring and incubated in ACSF supplemented with 5 µM 5-HT the “weak” TBS was ineffective in inducing a long-term change in FP amplitude, as in standard ACSF (con F vs. con F 5-HT: 101.6 ± 2.99%, n = 8 vs. 102.3 ± 4.96 mV, n = 8, respectively; q = 0.1644, *p* = 0.9994; Tukey’s multiple comparisons test; Fig. 2C1). However, in slices obtained from FLX-exposed females, supplementing ACSF with 5 µM 5-HT resulted in a significant reduction of FP amplitude after the “weak” TBS (FLX F vs. FLX F 5-HT: 99.52 ± 3.67%, n = 10 vs. 81.05 ± 4.14 mV, n = 10, respectively; q = 4.849, *p* = 0.0087; Tukey’s test; Fig. 2C2, C3).

### 3.4. Maternal fluoxetine alters the density of 5-HT innervation of the mPFC of female but not male offspring

As endogenous 5-HT might also contribute to LTD induction by the “weak” TBS protocol, we next compared the density of 5-HT fibers in the PL subdivision of mPFC in FLX-exposed and control male and female offspring mice. The analysis also involved infralimbic (IL) subdivision of mPFC as well as the motor cortex. Qualitative observation of 5-HT immunostaining (Fig. 3A) revealed some gross anatomical patterns in 5-HT innervation of three tested cortical areas (Fig. 3B). In particular, the outer cortical layers (I-III) consistently exhibited greater overall density of innervation than inner layers (V-VI). While all three anatomical sites were robustly innervated by 5-HT fibers with a similar overall pattern, the innervation within the outer layers of the motor cortex was comparatively the densest.

**Fig. 3.**
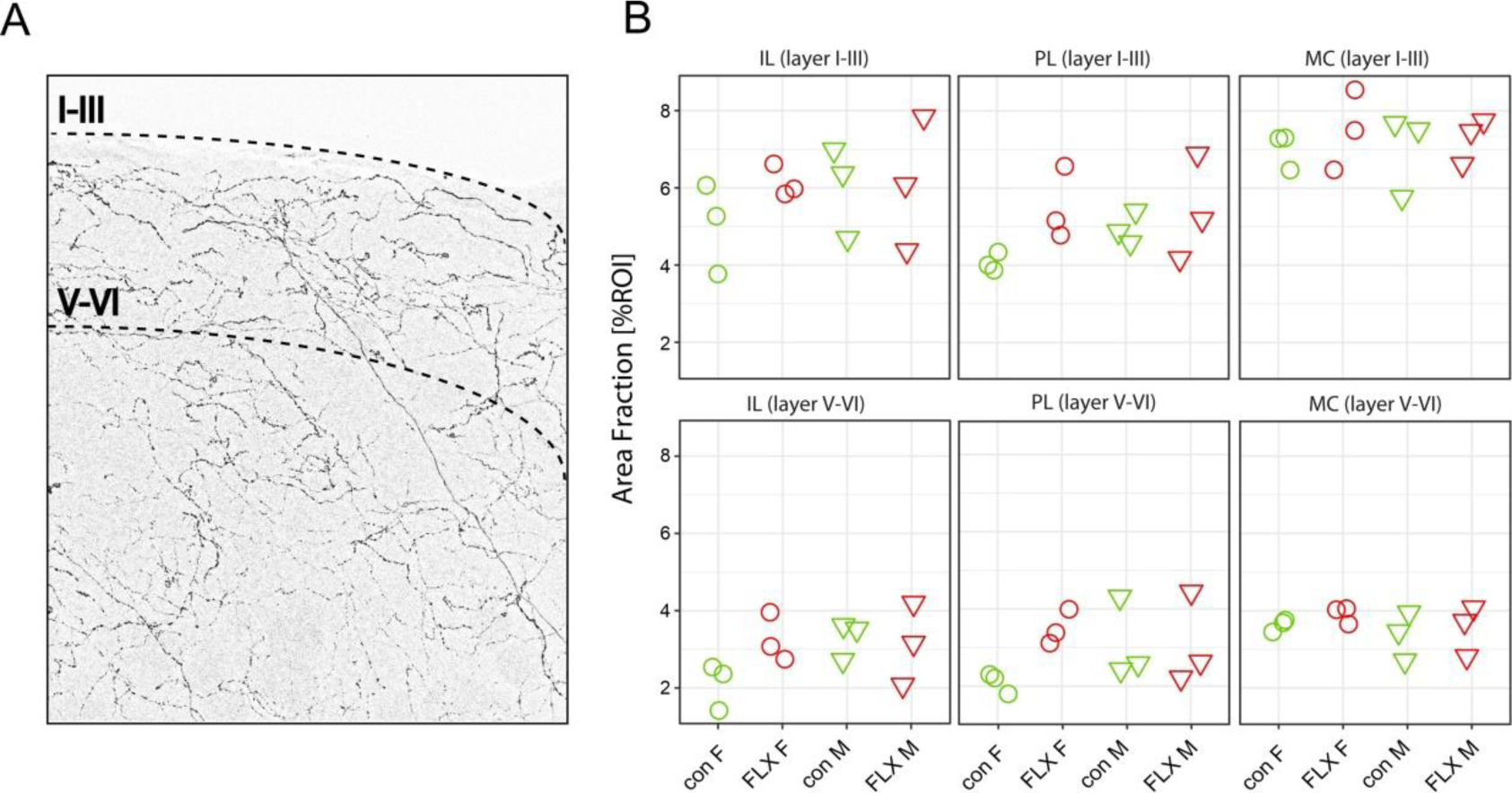
5-HT innervation of frontal cortical areas in FLX-exposed and control male and female offspring. (A) A representative microphotograph of 5-HT-immunopositive processes in the infralimbic cortex of a control female mouse. Dashed lines delineate areas of interest. Roman numerals indicate cortical layers. (B) Mean 5-HT-immunopositive area fraction (expressed as percentage of the total region of interest, %ROI) in respective experimental groups (n = 3 mice per group). IL – infralimbic, PL –prelimbic, MC – motor cortex. Note a greater density of innervation in superficial cortical layers (I-III) compared to inner layers (V-VI).

We then asked whether the density of serotonergic innervation of those cortical areas is altered by FLX treatment. For linear modeling, a full model (containing both variables and their interaction) was compared to a reduced model with only simple effects (no interaction), yielding a ΔAIC = 22.63, strongly in support of the simplified model (Tab. 1). Two-way ANOVA revealed significant effects of group (F(3.63) = 4.79, *p* < 0.005) as well as location (F(5,63) = 58.6, p < 0.0001). Of interest are the results of planned comparisons within group which reveal a lower baseline in the area fraction of 5-HT-positive processes in control females compared to males (con F vs. con M: ™0.59; confidence interval: ™1.12 to ™0.07, *p* < 0.05). This comparatively lower density of 5-HT innervation in control females was counteracted in the FLX-exposed group, which exhibited increased area fraction compared to controls (con F vs. FLX F: ™0.94; CI: ™1.46 to ™0.41, *p* < 0.001). Conversely, FLX treatment had no significant effects in males (con M vs. FLX M: ™0.16; CI: ™0.68 to 0.37, *p* > 0.5). While our analysis lacks sufficient power to distinguish between specific locations, data tentatively trend towards these differences being present in non-motor – i.e. medial prefrontal (PL and IL) – cortices, as opposed to motor cortex (Fig. 3B).

**Tab. 1.**
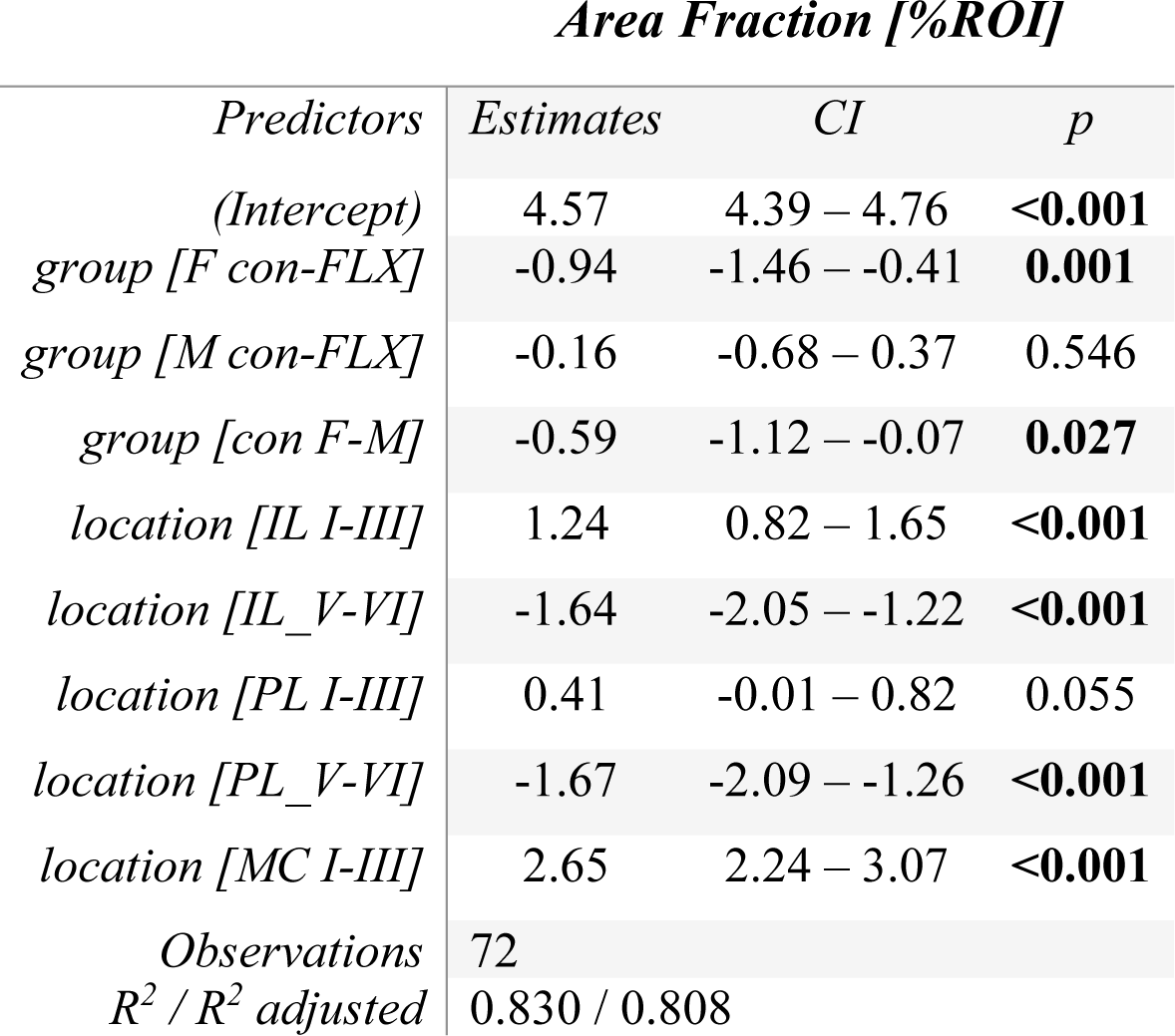
Model output with estimates, confidence intervals and corresponding *p*-values for levels of respective predictors (see: Material and methods for planned contrasts). Bolded *p*-values are < 0.05.

### 3.5. Lack of effect of maternal fluoxetine on neuronal excitability in the DRN of the offspring

The results obtained so far indicated that maternal exposure to FLX can potentially influence the density of 5-HT-positive processes in target cortical areas. In the next step we verified the influence of maternal FLX on electrophysiological properties of DRN 5-HT cells. Cells subjected to analysis demonstrated TPH2-immunoreactivity (Fig. 4A) and fired broad action potentials (duration: approx. 2 ms), characteristic of DRN 5-HT projection neurons (Fig. 4B1, B2; Galindo-Charles et al. 2008; Sowa et al. 2018). Two-way ANOVA revealed that there were no significant effects of FLX treatment, sex or their interaction on the resting membrane potential (main effect of FLX treatment: F(1, 70) = 0.001651, *p* = 0.9677; main effect of sex: F(1, 70) = 0.002888, *p* = 0.9573; interaction between FLX treatment and sex: F(1, 70) = 0.2763, *p* = 0.6008; Fig. 4C) and on input resistance (main effect of FLX treatment: F(1, 70) = 3.100, *p* = 0.0827; main effect of sex: F(1, 70) = 0.03829, *p* = 0.8454; FLX treatment x sex interaction: F(1, 70) = 0.1029, *p* = 0.7493; Fig. 4D). There were also no differences between FLX-exposed and control mice in the gain (main effect of FLX treatment: F(1, 70) = 1.949, *p* = 0.1671; main effect of sex: F(1, 70) = 0.07777, *p* = 0.7812; FLX treatment x sex interaction: F(1, 70) = 0.7873, *p* = 0.3780; Fig. 4E) and cumulative number of action potentials fired by DRN neurons upon testing (main effect of FLX treatment: F(1, 70) = 0.6673, *p* = 0.4168; main effect of sex: F(1, 70) = 2.004, *p* = 0.1613; FLX treatment x sex interaction: F(1, 70) = 1.941, *p* = 0.1679; Fig. 4F).

**Fig. 4.**
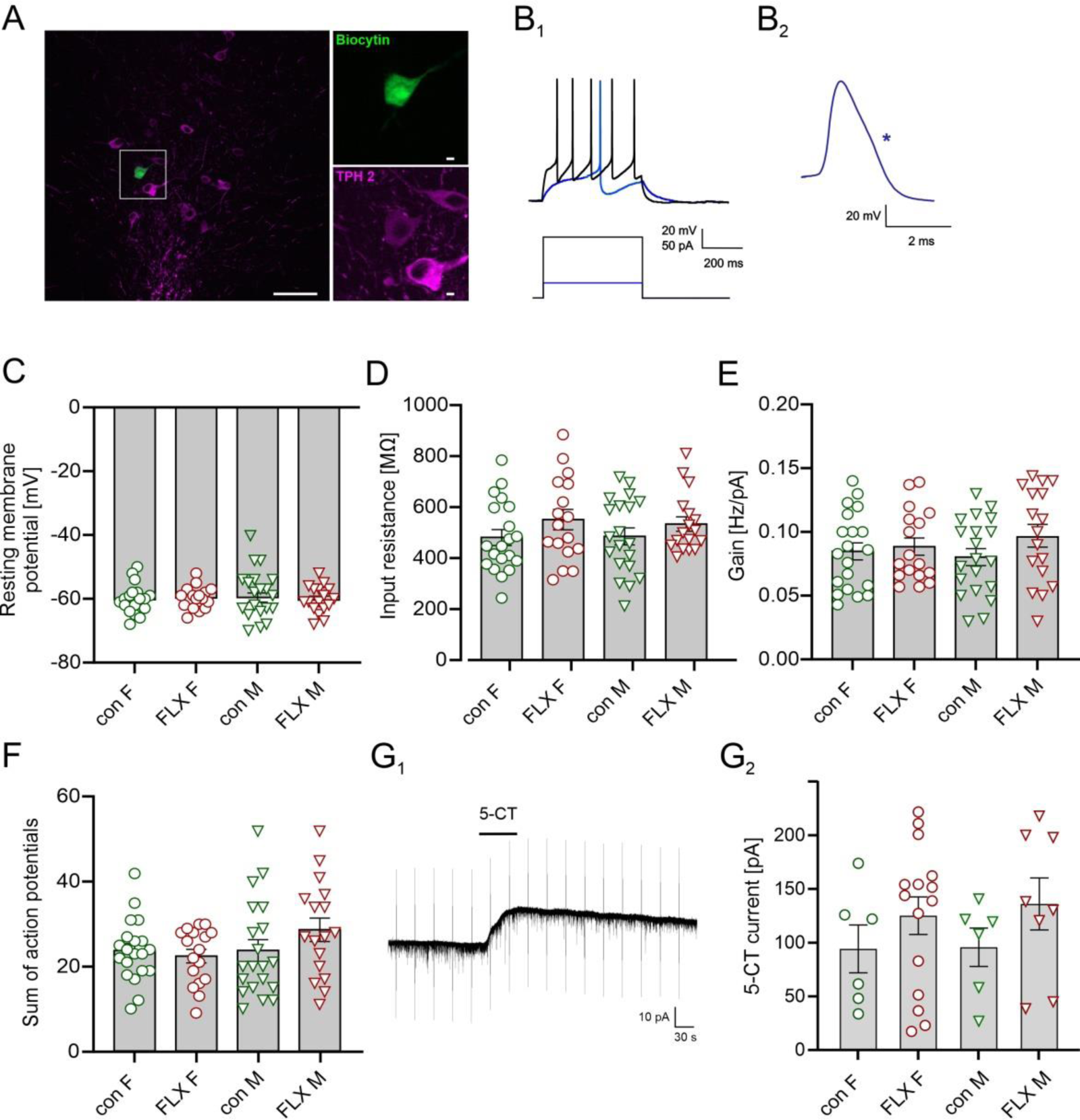
Maternal FLX exposure does not change membrane properties of DRN 5-HT neurons. (A) A microscope image of a coronal brain slice cut at the level of the DRN showing TPH2- immunopositive neurons (magenta) and a neuron filled with biocytin during whole-cell recording (green). Scale bar = 50 µm. Images to the right show an enlarged part of the image to the left (indicated by a square), confirming that the recorded cell was immunoreactive for TPH2. Scale bar = 5 µm. (B1) current-step stimulation at two intensities (bottom) and superimposed voltage responses of a representative DRN neuron (top) from a control female mouse. (B2) An action potential from a representative DRN neuron at expanded timescale with the “notch” on its descending phase marked with an asterisk. (C – F) summary graphs showing the resting membrane potential (C), input resistance (D), gain (E) and sum of action potentials (F) of recorded DRN neurons from four groups of mice. Shown are mean values ± SEM with circles and triangles representing individual neurons from female (F) and male (M) mice (con – green, FLX-exposed – red), respectively. (G1) current response of a representative DRN neuron from a control female mouse to the application of 5-CT (1 µM, horizontal bar). Vertical lines represent responses to test current pulses. (G2) summary graph showing the mean amplitude (± SEM) of 5-HT1A receptor-mediated current in four groups of mice. The differences between groups are not significant (two-way ANOVA, FLX exposure, sex, and FLX exposure x sex interaction: *p* > 0.05).

To assess the effect of the activation of 5-HT1A autoreceptors in DRN 5-HT neurons the outward current evoked by bath application of 5-CT (1 μM) was recorded (Fig. 4G1). There were no significant effects of FLX treatment and sex or their interaction on the amplitude of 5-CT induced current (main effect of FLX treatment: F(1, 27) = 1.294, *p* = 0.2652; main effect of sex: F(1, 27) = 0.1499, *p* = 0.7016; FLX treatment x sex interaction: F(1, 27) = 0.0298, *p* = 0.8642; two-way ANOVA; Fig. 4G2).

### 3.6. Maternal fluoxetine has little effect on synaptic transmission in the DRN of the offspring

To investigate whether maternal FLX exposure modifies spontaneous excitatory synaptic inputs to DRN neurons in the offspring, sEPSCs were recorded in whole-cell voltage clamp mode (Fig. 5A). While there was no significant effect of FLX treatment on sEPSC frequency (F(1, 41) = 0.2547, *p* = 0.6165), two-way ANOVA demonstrated an effect of sex (F(1, 41) = 1.91, *p* = 0.0013) and an interaction between FLX treatment and sex (F(1, 41) = 5.076, *p* = 0.0297). Mean sEPSC frequency was lower in DRN neurons in slices obtained from control female mice, compared to control males (con F vs. con M: 1.94 ± 0.34 Hz, n = 10 vs. 4.78 ± 0.65 Hz, n = 12, respectively; q = 5.633, *p* = 0.0015; Tukey’s multiple comparisons test; Fig. 5B). Two-way ANOVA also demonstrated a significant effect of sex on sEPSC amplitude (F(1, 41) = 7.866, *p* = 0.0077; Fig. 5C) but there was no significant effect of FLX treatment (F(1, 41) = 3.613, *p* = 0.0644) and no interaction between FLX treatment and sex (F(1, 41) = 2.408, *p* = 0.1284).

**Fig. 5.**
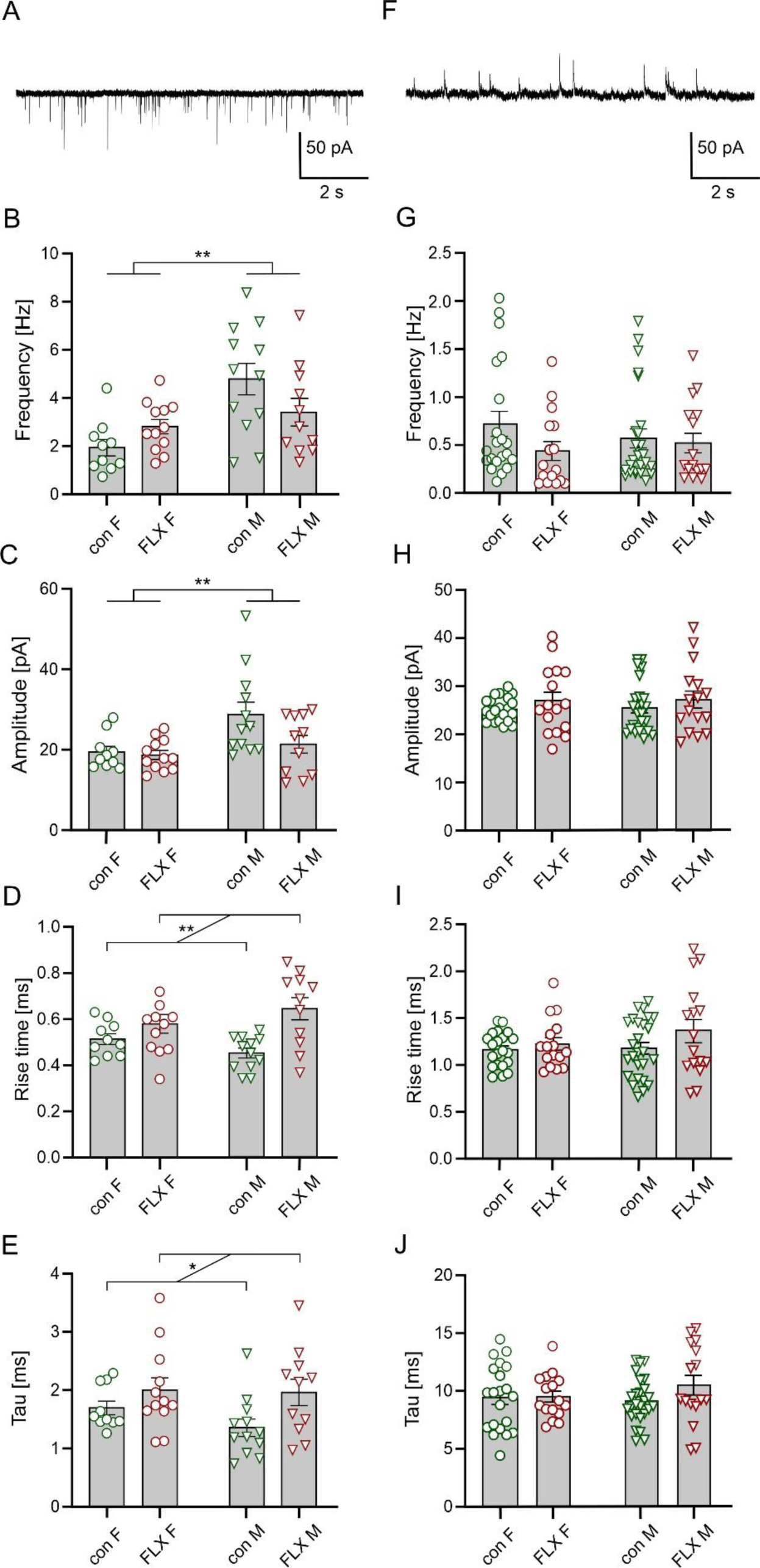
Excitatory and inhibitory synaptic transmission in DRN 5-HT neurons of FLX-exposed and control offspring. (A) Sample recording of sEPSCs from a representative DRN projection neuron of a control female mouse. (B) Summary graph showing sEPSC frequency in DRN projection neurons in four groups of mice. In this and the following graphs shown are mean values ± SEM with circles and triangles representing individual neurons from female (F) and male (M) mice (con – green, FLX-exposed – red), respectively. Two-way ANOVA, main effect of FLX exposure: *p* > 0.05, main effect of sex: \*\**p* < 0.01, FLX exposure x sex interaction: *p* < 0.05. (C) sEPSC amplitude in DRN projection cells. Two-way ANOVA, main effect of FLX exposure: *p* > 0.05, main effect of sex: \*\**p* < 0.01, FLX exposure x sex interaction: *p* > 0.05. (D) sEPSC rise time in DRN projection cells. Two-way ANOVA, main effect of FLX exposure: \*\**p* < 0.001, main effect of sex: *p* > 0.05, FLX exposure x sex interaction: *p* > 0.05. (E) sEPSC decay time constant in DRN projection cells. Two-way ANOVA, main effect of FLX exposure: \**p* < 0.05, main effect of sex: *p* > 0.05, FLX exposure x sex interaction: *p* > 0.05. (F) sample recording of sIPSCs from a representative DRN projection neuron. (G – J) summary graphs showing sIPSC frequency (G), amplitude (H), rise time (I) and decay time constant (J) in DRN projection cells in four groups of mice. In G-J differences between control and FLX-exposed male and female offspring are not significant (two-way ANOVA, FLX exposure, sex, and interaction of FLX exposure x sex: *p* > 0.05).

Two-way ANOVA revealed a significant effect of FLX exposure on parameters characterizing the kinetics of averaged sEPSCs waveforms: rise time (F(1, 41) = 13.48, *p* = 0.0007) and decay time constant (F(1, 41) = 6.126, *p* = 0.0175). There were, however, no significant effects of sex on either rise time (F(1, 41) = 0.0011, *p* = 0.9739) or decay time constant (F(1, 41) = 1.074, *p* = 0.3060) and there were also no interactions between FLX exposure and sex (rise time: F(1, 41) = 3.327, *p* = 0.0754); decay time constant: F(1, 41) = 0.6572, *p* = 0.4222). Mean (±SEM) rise times of averaged sEPSC were as follows - con F: 0.51 ± 0.02 ms, n = 10; FLX F: 0.58 ± 0.04 ms, n = 12; con M: 0.45 ± 0.02 ms, n = 12; FLX M: 0.64 ± 0.05 ms, n = 11. Mean decay time constants of averaged sEPSC were as follows - con F: 1.69 ± 0.11 ms, n = 10; FLX F: 2.0 ± 0.20 ms, n = 12; con M: 1.35 ± 0.14 ms, n = 12; FLX M: 1.96 ± 0.22 ms, n = 11.

Two-way ANOVA of the mean frequency, amplitude, rise time and decay time constant of sIPSCs recorded from DRN neurons (Fig. 5F) did not reveal any significant differences between control and FLX-exposed male and female offspring (Fig 5G-J).

### 3.7. Maternal fluoxetine alters the morphology of DRN 5-HT neurons of female but not male offspring

To assess if maternal FLX exposure affects the morphology of DRN 5-HT neurons, 3D reconstructions of the axodendritic tree of cells filled with biocytin during whole-cell recording (Fig. 6A) were subjected to Sholl analysis. Sholl profiles were next averaged, and their polynomial fits were compared using the AIC metric (Fig. 6B, C) where a full model (separate polynomial fits for control/FLX groups) was compared to a reduced model with only one fit for combined datasets, yielding a ΔAIC = 60.57, strongly in favor of the full model. In contrast, comparison of fits representing Sholl profiles of cells obtained from FLX- exposed and control males revealed a ΔAIC of ™4.292), which was in favor of the reduced model.

**Fig. 6.**
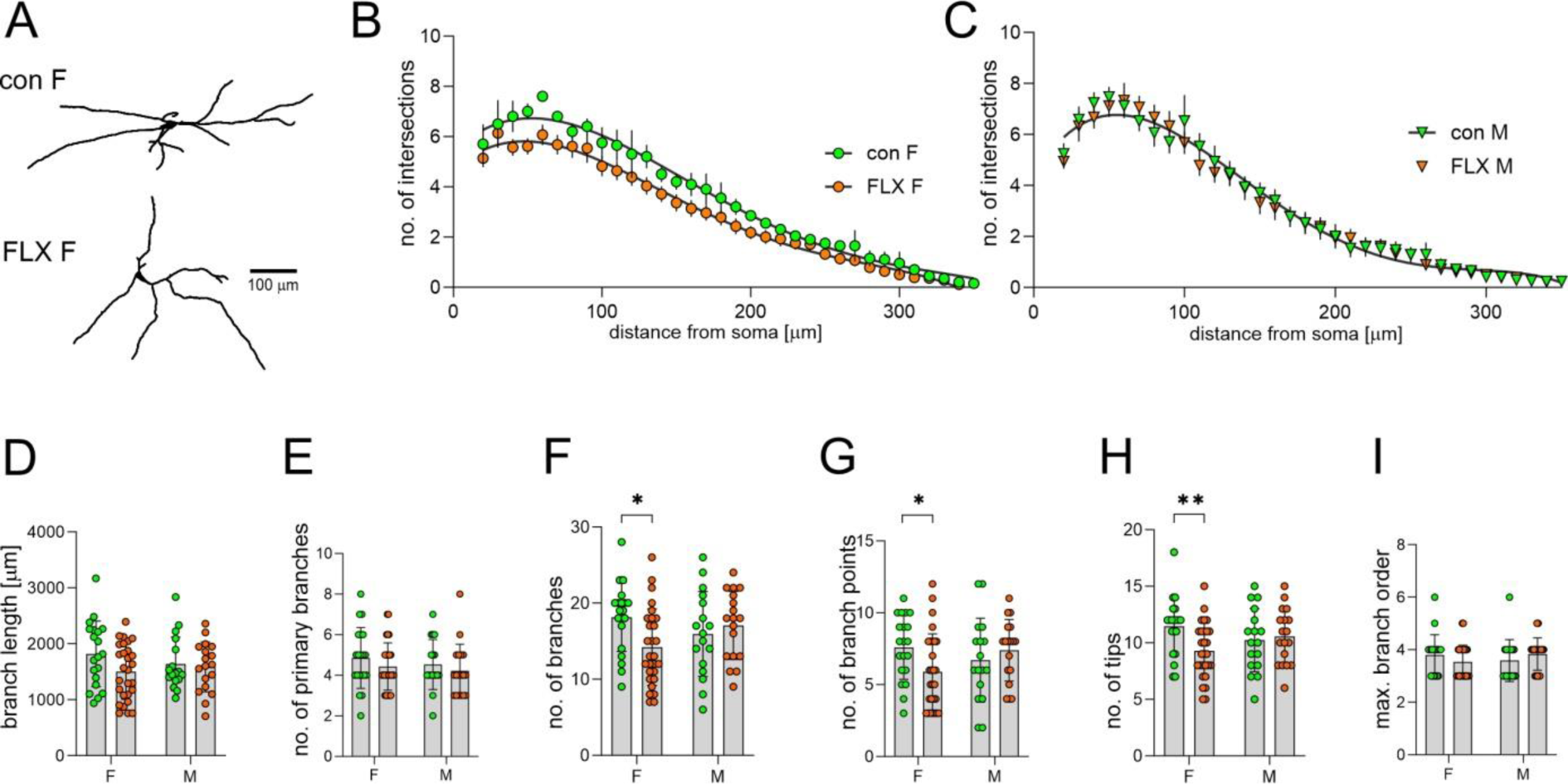
Maternal FLX exposure decreases the axodendritic tree complexity of DRN 5-HT neurons of female but not male offspring. (A) 3D tracings of representative DRN 5-HT neurons originating from FLX-exposed (F FLX) and control (con F) female offspring. (B) Sholl profiles of 5-HT neurons of female DRN. Graphs show the mean number (± SEM) of intersections of Sholl circles with cells’ neurites (con – green, FLX-exposed – red) with black lines representing the polynomial fits to data. The AIC metric indicated a significant difference between the two fits. (C) Sholl profiles of 5-HT cells of male DRN (con – green, FLX-exposed – red) with black lines representing the polynomial fit to data. The AIC metric indicated that one fit represents both profiles. (D – I) summary graphs showing the total branch length (D), number of primary branches (E), total number of branches (F), number of branch points (G), number of branch tips (H), and maximum branch order (I) in DRN cells from four groups of mice. Two-way ANOVA, multiple comparisons: \**p* < 0.05, \*\**p* < 0.01.

Amongst morphological features describing the morphology of DRN neurons that were measured, there was a significant interaction between FLX treatment and sex regarding the total number of branches (two-way ANOVA; F(1, 79) = 5.28, *p* = 0.0242; Fig. 6F), number of branch points (F(1, 79) = 4.536, *p* = 0.0363; Fig. 6G), and number of branch tips (F(1, 79) = 4.722, *p* = 0.038; Fig. 6H) that were significantly smaller (Sidak’s multiple comparisons test, *p* < 0.05) in FLX-exposed female but not in male offspring, compared to control. There were no significant effects of either FLX treatment or sex regarding these parameters. On the other hand, there were no effects of sex, FLX exposure and no interaction between FLX exposure and sex (two-way ANOVA, *p* > 0.05), regarding the total branch length (Fig. 6D), number of primary branches (Fig. 6E), and maximum branch order (Fig. 6I).

### 3.8. Maternal vortioxetine does not affect field potentials and long-term potentiation in the mPFC of female and male offspring

In the final set of experiments, we tested whether a relatively new antidepressant, vortioxetine, affects synaptic transmission and long-term plasticity in layer II/III of the PL subdivision of mPFC in a similar way to FLX. There were no significant effects of VOR treatment, sex or their interaction on the maximum amplitude of FPs recorded under basal conditions, calculated using Boltzmann fits (two-way ANOVA, main effect of VOR treatment: F(1, 29) = 1.839, *p* = 0.1855, main effect of sex: F(1, 29) = 1.839, *p* = 0.1839, interaction between treatment and sex: F(1, 29) = 0.0025, *p* = 0.9606; Fig. 7A-C).

**Fig. 7.**
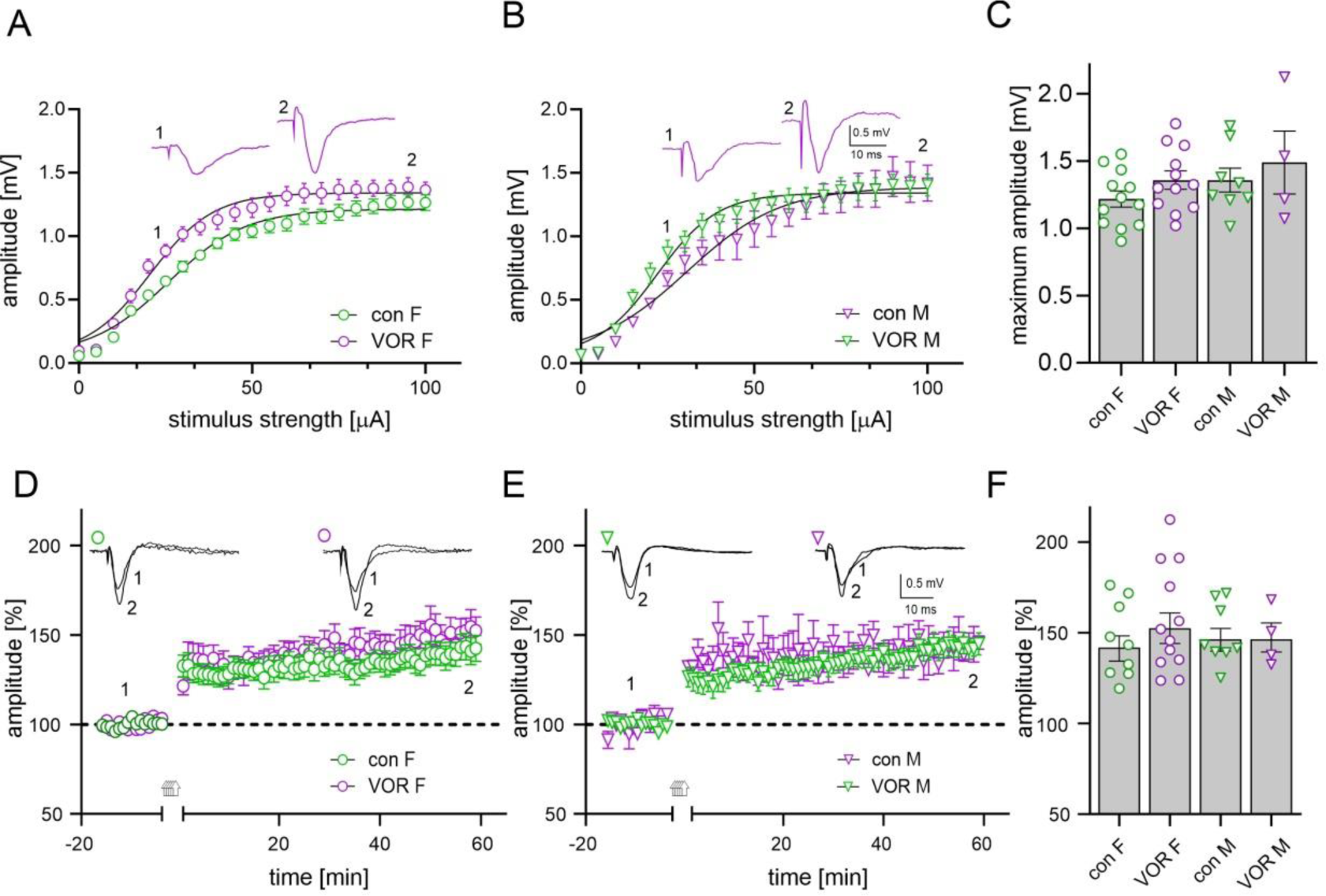
Maternal vortioxetine affects neither FPs nor LTP in the mPFC of the female and male offspring. Graphs (A) and (B) illustrate the effect of vortioxetine (VOR) exposure on the relationship between stimulus intensity and mean FP amplitude (± SEM) in mPFC slices obtained from female (A) and male (B) offspring that had been exposed to VOR (magenta) and control mice (con, green). Insets in A and B show representative FPs recorded at two stimulation intensities (1, 2). Black lines represent fits to the Boltzmann equation. (C) Comparison of calculated maximum FP amplitudes in four groups of animals. Shown are mean values ± SEM with circles and triangles representing individual slices. Two-way ANOVA - VOR exposure, sex, and VOR exposure x sex interaction: *p* > 0.05). Symbols and labels as in A, B. (D, E) Plots of the amplitude of FPs (mean ± SEM) in slices obtained from control (con F, green) and VOR-exposed females (VOR F, magenta; D) as well as from control (con M, green) and VOR-exposed males (VOR M, magenta; E). Arrows denote the time of theta-burst stimulation (TBS, repeated 5 times). Insets in D and E show superpositions of FPs recorded in the course of representative experiments before and after TBS at times indicated by numbers. (F) Mean (± SEM) amplitude of FPs recorded between 45 - 60 min after TBS, relative to baseline. Two-way ANOVA, VOR exposure, sex, and VOR exposure x sex interaction: *p* > 0.05).

Similarly, two-way ANOVA did not reveal any significant effect of VOR exposure (F(1, 29) = 0.0085, *p* = 0.9274), sex (F(1, 29) = 0.3985, *p* = 0.5328), and interaction between VOR exposure and sex (F(1, 29) = 0.3985, *p* = 0.5328) on the amplitude of FPs recorded between 45 - 60 min after stimulation of slices with the standard TBS protocol (5 trains of TBS) relative to baseline. Thus, we conclude that in contrast to FLX, maternal exposure to VOR does not exert detrimental effects on glutamatergic transmission and LTP in mPFC of the offspring.

## 4. Discussion

The consequences of maternal FLX exposure on behavior of the offspring in later life have previously been investigated extensively, however, reported effects are divergent possibly due to variability of the duration of exposure to the drug during pre-and postnatal development and different doses of FLX administered (reviewed in: Glover and Clinton, 2016; Ramsteijn et al., 2020). Only a few employed exposure of dams to FLX during the whole pregnancy and lactation period, an approach that appears to be the most relevant to the timing of maternal FLX exposition in humans. These studies reported e.a. occurrence of anxiety-and depression-like phenotypes in the offspring as well as occurrence of repetitive patterns of behavior (Kiryanova et al., 2013; Kinast et al., 2013; Maloney et al., 2018). In line with Lisboa et al. (2007), who reported that exposure of mice dams to FLX throughout gestation and lactation results in an increase in immobility time in the forced swimming test in female offspring, we found that FLX-exposed female mice demonstrated decreased sucrose preference, indicative of anhedonia (Primo et al., 2023). Based on Lisboa et al. (2007), in our study dams received FLX with the drinking water in a dose of approx. 7.5 mg/kg/day.

Experiments on rats confirmed that maternal exposure to FLX in a slightly higher dose (10 mg/kg/day) from GD 0 to PND 14 induced anxiety-like and depressive-like phenotype in male and female offspring (Millard et al., 2019; 2021). To test the stereotypic behavior patterns of the offspring, we performed the marble burying test as a measure of repetitive and perseverative behavior (Thomas et al., 2009) but we observed no difference between FLX-exposed and control female offspring in this task. However, FLX-exposed females demonstrated disturbance of the temporal order memory, a capability that depends on proper functioning of the mPFC (Hannesson et al., 2004; DeVito and Eichenbaum, 2011), that coincided with a reduced amplitude of FPs in the PL subdivision in the mPFC.

Since short-latency FPs in rodent mPFC are a product of the activity of monosynaptic glutamatergic connections (Wallace et al., 2014), decreased amplitude of FPs in slices originating from FLX-exposed mice is suggestive of a reduction of the excitatory transmission in the mPFC. Importantly, we observed these effects in female but not in male offspring. In rats, the first 5-HT neurons appear on GD 12 and 5-HTergic afferents reach the telencephalon by GD 15 (Lauder, 1990). 5-HT receptors, transporters and catabolic enzymes are expressed concurrently and the innervation pattern of target structures by 5-HTergic fibers develops throughout the embryonic and postnatal life until PND 21 (Lauder, 1990). The level of 5-HT in the developing rodent brain peaks within the first postnatal week, after which it declines, reaching adult level at PND 15 (Hohmann et al., 1988). Thus, maternal FLX and resulting, higher than normal 5-HT level in the developing brain may profoundly distort early and late phases of the development of brain neural circuitry. The mechanism of the observed reduction of the excitatory transmission in the mPFC remains to be established but we note that no changes in the NMDA or AMPA receptor subunit expression in the mPFC in female adolescent offspring of Sprague-Dawley rats were observed after maternal FLX exposure (Millard et al., 2021). Interestingly, perinatal FLX exposure reduced GluN1, GluN2A and PSD-95 levels in rat male offspring (Millard et al., 2019). Augmented inhibitory synaptic transmission due to increased excitability of GABAergic interneurons has been reported to occur in layer V mPFC neurons of male mice after maternal treatment with FLX, however, in that study FLX was administered prenatally (GD 4 - GD 19; Yu et al., 2019). The discrepancies between the results of these studies and ours are likely to result from species and experimental design differences. In a study employing a similar methodology to ours, maternal FLX exposure throughout gestation and lactation was found to decrease dendritic complexity and spine density of layer V mPFC pyramidal neurons in young adult offspring mice (Maloney et al., 2022). Thus, our findings on the function of neuronal circuitry in mPFC of FLX-exposed mice complement earlier observations on mouse mPFC neuronal morphology. One of the consequences of weakened excitatory transmission in the mPFC of female but not male offspring appears to be an impairment of LTP.

Importantly, the results of the present study indicate that feeding of dams with VOR- containing food at the dose that produces procognitive effects (Felice et al., 2018) and reverses cognitive impairments (Pehrson et al., 2018) does not result in adverse effects on excitatory synaptic transmission in offspring mPFC, in contrast to FLX. Besides, our recordings of layer II/III FPs in control mice do not confirm the notion that females express a higher level of glutamatergic transmission in the mPFC than males (Knouse et al., 2022), which was also based on recordings from layer V neurons.

It has recently been shown that female offspring of rats’ dams that were treated during pregnancy and lactation with another SSRI, citalopram, exhibited increased expression of mRNAs for 5-HT1B, 5-HT2A and 5-HT2C receptors in the mPFC (Unroe et al., 2022). Postnatal exposure to citalopram resulted in reduced expression of 5-HT transporter in rat cortex (Maciag et al., 2006). Since 5-HT receptors modulate glutamatergic transmission and neuronal membrane excitability (Higa et al., 2022; Zhong et al., 2008a, b), if similar effects occur in mice exposed to FLX, they might contribute to alterations in the induction of long-term synaptic plasticity observed in our experiments. In earlier studies aimed at finding facilitatory effects of 5-HT on the induction of LTD in mPFC slices, a higher concentration of 5-HT was used (40-50 μM), which markedly reduced evoked responses upon application (Higa et al., 2022; Zhong et al., 2008a, b). We used a one order of magnitude lower ambient concentration of 5-HT that by itself did not affect recorded responses but, nevertheless, it enabled LTD induction in slices originating from FLX-exposed mice. This effect coincided with indications of increased density of 5-HT-immunoreactive fibers in the mPFC but not the motor cortex of FLX-exposed female offspring. It has been reported that prenatal FLX exposure between GDs 13-20 results in a decrease in 5-HT content in the frontal cortex of adult rat offspring (Cabrera-Vera et al., 1997) but again, the discrepancy between the results of that study and ours is likely to result from species and experimental design differences. In our experiments, in control mice, subthreshold TBS stimulation protocol combined with the application of 5 μM 5-HT proved to be ineffective for LTD induction. It is tempting to speculate that in FLX-exposed female offspring the combination of the same 5-HT bath concentration with a larger amount of 5-HT released from denser 5-HT fibers upon TBS stimulation of the tissue and altered expression of 5-HT receptors allows for crossing the LTD induction threshold upon application of the “weak” TBS protocol.

In addition to a possible increase of 5-HT fiber density in the mPFC, the results of the present study show that maternal FLX reduces the axodendritic tree complexity of DRN 5-HT neurons in female but not male offspring. While in our study dams received FLX during pregnancy and lactation, earlier studies aimed at finding the effects of maternal FLX on DRN neurons tested the effects of the drug delivered in shorter periods. In rats exposed to FLX between PNDs 1-21 a decreased number of 5-HT neurons having a smaller size of cell bodies was observed in the DRN (Silva et al., 2010). Postnatal exposure to citalopram resulted in a reduced TPH expression in the DRN (Maciag et al., 2006). DRN 5-HT neurons undergo marked changes in morphology during development, especially during the first three postnatal weeks (Rood et al., 2014), and our data indicate that these processes are distorted by maternal FLX, at least in female offspring.

However, these structural modifications are not accompanied by modifications in electrophysiological properties of DRN 5-HT neurons as neither intrinsic excitability nor 5- HT1A receptor-mediated current were significantly altered in the DRN of FLX-exposed offspring of either sex. A lack of alteration in 5-HT1A receptor-mediated current is consistent with a lack of the effect of FLX exposure on performance in the marble burying test as the latter is sensitive to activation of presynaptic 5-HT1A receptors (Depoortere et al., 2021). While no effects of maternal FLX on inhibitory postsynaptic currents (sIPSCs) in DRN 5-HT neurons were evident, we observed FLX exposure-related increases in the values of parameters characterizing the kinetics of averaged sEPSCs waveforms: rise time and decay time constant. Elucidation of the mechanism underlying this effect will require further studies, but we note that it might, potentially, involve altered cable filtering of the neuronal membrane (Kleppe and Robinson, 1999) or result from an increased proportion of excitatory synapses located at a larger distance from the soma. It has been reported that decay time constant of sEPSCs recorded from DRN 5-HT neurons changes over development (Kisner and Polter, 2023) and exposure to FLX from PND 2 to PND 14 increases the number of synapses on DRN neurons that are formed by the glutamatergic projection from the mPFC but not from other sources (Soiza-Reilly et al., 2019).

## 5. Conclusion

In the present study we demonstrate that exposure of mouse dams to FLX during gestation and lactation decreases glutamate-mediated FPs, impairs LTP and facilitates induction of LTD in the mPFC of adolescent offspring. Notably, these effects occur in female but not male mice. FLX-exposed female offspring express poor performance in the temporal order memory task and reduced sucrose preference with no change in marble burying behavior. Maternal FLX tends to increase the density of 5-HTergic fibers in the mPFC and reduces the axodendritic tree complexity of DRN 5-HT neurons in female but not male offspring, however, these structural modifications are not accompanied by significant alterations in the excitability of DRN 5-HT cells. There is a mild effect of FLX treatment on excitatory synaptic inputs to DRN neurons of either sex, but inhibitory inputs remain unchanged. The results of the present study also indicate that neither FPs nor LTP are impaired in the mPFC of the offspring after maternal exposure to vortioxetine.

### CRediT authorship contribution statement

**Bartosz Bobula:** Methodology, Formal Analysis, Investigation,Visualization, Resources. **Joanna Bąk:** Formal Analysis, Investigation, Visualization**. Agnieszka Kania:** Formal Analysis, Investigation. **Marcin Siwiec:** Conceptualization, Formal Analysis, Investigation, Writing – Review & Editing**. Michał Kiełbiński:** Formal Analysis, Investigation, Writing – Review & Editing. **Krzysztof Tokarski:** Conceptualization, Visualization. **Agnieszka Pałucha-Poniewiera:** Conceptualization. Investigation. **Grzegorz Hess:** Conceptualization, Methodology, Formal analysis, Investigation, Writing – Original Draft, Supervision, Funding acquisition

### Data availability

Data will be made available on request.

## Acknowledgements

This work was supported by the National Science Center, Poland, grant DEC- 2017/27/B/NZ4/01527.

## References

1. Adjimann, T.S., Argañaraz, C.V., Soiza-Reilly, M., 2021. Serotonin-related rodent models of early-life exposure relevant for neurodevelopmental vulnerability to psychiatric disorders. Transl. Psychiatry 11, 280. 10.1038/s41398-021-01388-6.

2. Andalib, S., Emamhadi, M.R., Yousefzadeh-Chabok, S., Shakouri, S.K., Høilund-Carlsen, P.F. et al., 2017. Maternal SSRI exposure increases the risk of autistic offspring: A meta-analysis and systematic review. Eur. Psychiatry 45, 161–166. 10.1016/j.eurpsy.2017.06.001.

3. Beck, S.G., Pan, Y.-Z., Akanwa, A.C., Kirby, L.G. (2004) Median and dorsal raphe neurons are not electrophysiologically identical. J Neurophysiol 91, 994–1005. 10.1152/jn.00744.2003.

4. Besag, F.M.C., Vasey, M.J., 2023. Should Antidepressants be Avoided in Pregnancy? Drug Saf. 46, 1–17. 10.1007/s40264-022-01257-1.

5. Cabrera-Vera, T.M., Garcia, F., Pinto, W., Battaglia, G., 1997. Effect of prenatal fluoxetine (Prozac) exposure on brain serotonin neurons in prepubescent and adult male rat offspring. J. Pharmacol. Exp. Ther. 280, 138–145.

6. Chandler, D.J., Lamperski, C.S., Waterhouse, B.D., 2013. Identification and distribution of projections from monoaminergic and cholinergic nuclei to functionally differentiated subregions of prefrontal cortex. Brain Res. 1522, 38–58. 10.1016/j.brainres.2013.04.057.

7. Deacon, R.M.J., 2006. Digging and marble burying in mice: simple methods for in vivo identification of biological impacts. Nat. Protoc. 1, 122–124. 10.1038/nprot.2006.20.

8. Depoortere, R., Bardin, L., Auclair, A.L., Bruins Slot, L.A., Newman-Tancredi, A., 2021. Marble burying in NMRI male mice is preferentially sensitive to pre-versus postsynaptic 5-HT1A receptor biased agonists. Pharmacology 106, 114–118. doi: 10.1159/000509729.

9. Desaunay, P., Eude, L.-G., Dreyfus, M., Alexandre, C., Fedrizzi, S., et al., 2023. Benefits and risks of antidepressant drugs during pregnancy: a systematic review of meta-analyses. Paediatr. Drugs 25:247–265. 10.1007/s40272-023-00561-2.

10. DeVito, L.M., Eichenbaum, H., 2011. Memory for the order of events in specific sequences: contributions of the hippocampus and medial prefrontal cortex. J. Neurosci. 31, 3169 – 3175. doi: 10.1523/JNEUROSCI.4202-10.2011.

11. Domingues, R.D., Wiltbank, M.C., Hernandez, L.L., 2023. The antidepressant fluoxetine (Prozac®) modulates estrogen signaling in the uterus and alters estrous cycles in mice. Mol. Cell. Endocrinol. 559, 111783. 10.1016/j.mce.2022.111783.

12. Felice, D., Guilloux, J.-P., Pehrson, A., Li, Y., Mendez-David, I., Gardier, A.M., Sanchez, C., David, D.J., 2018. Vortioxetine improves context discrimination in mice through a neurogenesis independent mechanism. Front. Pharmacol. 9, 204. 10.3389/fphar.2018.00204.

13. Galindo-Charles, L., Hernandez-Lopez, S., Galarraga, E., Tapia, D., Bargas, J., Garduño, J., Frías-Dominguez, C., Drucker-Colin, R., Mihailescu, S., 2008. Serotoninergic dorsal raphe neurons possess functional postsynaptic nicotinic acetylcholine receptors. Synapse 62, 601–615. 10.1002/syn.20526.

14. Glover, M.E., Clinton, S.M., 2016. Of rodents and humans: A comparative review of the neurobehavioral effects of early life SSRI exposure in preclinical and clinical research. Int. J. Dev. Neurosci. 51, 50–72. 10.1016/j.ijdevneu.2016.04.008.

15. Hannesson, D.K., Vacca, G., Howland, J.G., Phillips, A.G., 2004. Medial prefrontal cortex is involved in spatial temporal order memory but not spatial recognition memory in tests relying on spontaneous exploration in rats. Behav. Brain Res. 153, 273–285. 10.1016/j.bbr.2003.12.004.

16. Hayes, R.M., Wu, P., Shelton, R.C., Cooper, W.O., Dupont, W.D., Mitchel, E., Hartert, T.V., 2012. Maternal antidepressant use and adverse outcomes: a cohort study of 228,876 pregnancies. Am. J. Obstet. Gynecol. 207, 49.e1–9. 10.1016/j.ajog.2012.04.028.

17. Heikkinen, T., Ekblad, U., Palo, P., Laine, K., 2003. Pharmacokinetics of fluoxetine and norfluoxetine in pregnancy and lactation. Clin. Pharmacol. Ther. 73, 330–337. 10.1016/s0009-9236(02)17634-x.

18. Hiemke, C., Härtter, S., 2000. Pharmacokinetics of selective serotonin reuptake inhibitors. Pharmacol. Ther. 85, 11–28. 10.1016/s0163-7258(99)00048-0.

19. Higa, G.S.V., Francis-Oliveira, J., Carlos-Lima, E., Tamais, A.M., Borges, F.D.S., Kihara, A.H., Shieh, I.C., Ulrich, H., Chiavegatto, S., De Pasquale, R., 2022. 5-HT-dependent synaptic plasticity of the prefrontal cortex in postnatal development. Sci. Rep.12, 21015. 10.1038/s41598-022-23767-9.

20. Hohmann, C.F., Hamon, R., Batshaw, M.L., Coyle, J.T., 1988. Transient postnatal elevation of serotonin levels in mouse neocortex. Brain Res. 471, 163–166. 10.1016/0165-3806(88)90163-0.

21. Jacobs, B.L., Azmitia, E.C., 1992 Structure and function of the brain serotonin system. Physiol. Rev. 72, 165–229. 10.1152/physrev.1992.72.1.165.

22. Kim, J., Riggs, K.W., Misri, S., Kent, N., Oberlander, T.F., et al., 2006. Stereoselective disposition of fluoxetine and norfluoxetine during pregnancy and breast-feeding. Br. J. Clin. Pharmacol. 61, 155–163. 10.1111/j.1365-2125.2005.02538.x.

23. Kinast, K., Peeters, D., Kolk, S.M., Schubert, D., Homberg, J.R., 2013. Genetic and pharmacological manipulations of the serotonergic system in early life: neurodevelopmental underpinnings of autism-related behavior. Front. Cell. Neurosci. 7, 72. 10.3389/fncel.2013.00072.

24. Kiryanova, V., McAllister, B.B., Dyck, R.H., 2013. Long-term outcomes of developmental exposure to fluoxetine: a review of the animal literature. Dev. Neurosci. 35, 437–439. 10.1159/000355709.

25. Kiryanova, V., Meunier, S.J., Vecchiarelli, H.A., Hill, M.N., Dyck, R.H., 2016. Effects of maternal stress and perinatal fluoxetine exposure on behavioral outcomes of adult male offspring. Neuroscience 320, 281–296. 10.1016/j.neuroscience.2016.01.064.

26. Kisner, A., Polter, A.M., 2023.Maturation of glutamatergic transmission onto dorsal raphe serotonergic neurons. bioRxiv 2023.01.19.524776. doi: 10.1101/2023.01.19.524776. Preprint.

27. Kleppe, I.C., Robinson, H.P., 1999. Determining the activation time course of synaptic AMPA receptors from openings of colocalized NMDA receptors. Biophys. J. 77, 1418– 1427. doi: 10.1016/S0006-3495(99)76990-0.

28. Knouse, M.C., McGrath, A.G., Deutschmann, A.U., Rich, M.T., Zallar, L.J., Rajadhyaksha, A.M., Briand, L.A., 2022. Sex differences in the medial prefrontal cortical glutamate system. Biol. Sex Differ. 13, 66. 10.1186/s13293-022-00468-6.

29. Kolk, S.M., Rakic, P., 2022. Development of prefrontal cortex. Neuropsychopharmacol. 47, 41–57. 10.1038/s41386-021-01137-9.

30. Lauder, J.M., 1990. Ontogeny of the serotonergic system in the rat: serotonin as a developmental signal. Ann. N. Y. Acad. Sci. 600, 297–313. 10.1111/j.1749-6632.1990.tb16891.x.

31. Lebin, L.G., Novick, A.M., 2022. Selective Serotonin Reuptake Inhibitors (SSRIs) in pregnancy: an updated review on risks to mother, fetus, and child. Curr. Psychiatry Rep. 24, 687–95. 10.1007/s11920-022-01372-x.

32. Lisboa, S.F., Oliveira, P.E., Costa, L.C., Venâncio, E.J., Moreira, E.G., 2007. Behavioral evaluation of male and female mice pups exposed to fluoxetine during pregnancy and lactation. Pharmacology 80, 49–56. 10.1159/000103097.

33. Longair, M.H., Baker, D.A., Armstrong, J.D., 2011. Bioinformatics 27, 2453–2454. 10.1093/bioinformatics/btr390.

34. Maciag, D., Simpson, K.L., Coppinger, D., Lu, Y., Wang, Y., et al., 2006. Neonatal antidepressant exposure has lasting effects on behavior and serotonin circuitry. Neuropsychopharmacology 31, 47–57. 10.1038/sj.npp.1300823.

35. Maloney, S.E., Akula, S., Rieger, M.A., McCullough, K.B., Chandler, K., Corbett, A.M., McGowin, A.E., Dougherty, J.D., 2018. Examining the reversibility of long-term behavioral disruptions in progeny of maternal SSRI exposure. eNeuro 5, ENEURO.0120- 18.2018. 10.1523/ENEURO.0120-18.2018.

36. Maloney, S.E., Tabachnick, D.R., Jakes, C., Avdagic, S., Bauernfeind, A.L., Dougherty, J.D., 2022. Fluoxetine exposure throughout neurodevelopment differentially influences basilar dendritic morphology in the motor and prefrontal cortices. Sci. Rep. 12, 7605. 10.1038/s41598-022-11614-w.

37. McAllister, B.B., Kiryanova, V., Dyck, R.H., 2012. Behavioural outcomes of perinatal maternal fluoxetine treatment. Neuroscience 226, 356–366. 10.1016/j.neuroscience.2012.09.024.

38. Millard, S.J., Lum, J.S., Fernandez, F., Weston-Green, K., Newell, K.A., 2019. Perinatal exposure to fluoxetine increases anxiety-and depressive-like behaviours and alters glutamatergic markers in the prefrontal cortex and hippocampus of male adolescent rats: A comparison between Sprague-Dawley rats and the Wistar-Kyoto rat model of depression. J. Psychopharmacol. 33, 230–243. 10.1177/0269881118822141.

39. Millard, S.J., Lum, J.S., Fernandez, F., Weston-Green, K., Newell, K.A., 2021. The effects of perinatal fluoxetine exposure on emotionality behaviours and cortical and hippocampal glutamatergic receptors in female Sprague-Dawley and Wistar-Kyoto rats. Prog. Neuropsychopharmacol. Biol. Psychiatry 108, 110174. 10.1016/j.pnpbp.2020.110174.

40. Molenaar, N.M., Bais, B., Lambregtse-van den Berg, M.P., Mulder, C.L., Howell, E.A., et al., 2020. The international prevalence of antidepressant use before, during, and after pregnancy: A systematic review and meta-analysis of timing, type of prescriptions and geographical variability. J. Affect. Disord. 264, 82–89. 10.1016/j.jad.2019.12.014.

41. Muzerelle, A., Scotto-Lomassese, S., Bernard, J.F., Soiza-Reilly, M., Gaspar, P., 2016. Conditional anterograde tracing reveals distinct targeting of individual serotonin cell groups (B5-B9) to the forebrain and brainstem. Brain Struct. Funct. 221, 535–561. 10.1007/s00429-014-0924-4.

42. Navailles, S., Hof, P.R., Schmauss, C., 2008. Antidepressant drug-induced stimulation of mouse hippocampal neurogenesis is age-dependent and altered by early life stress. J. Comp. Neurol. 509, 372–381. 10.1002/cne.21775.

43. Oh, J., Zupan, B., Gross, S., Toth, M., 2009. Paradoxical anxiogenic response of juvenile mice to fluoxetine. Neuropsychopharmacology 34, 2197–2207. 10.1038/npp.2009.47.

44. Pałucha-Poniewiera, A., Bobula, B., Rafało-Ulińska, A., 2023. The antidepressant-like activity and cognitive enhancing effects of the combined administration of (R)-ketamine and LY341495 in the CUMS model of depression in mice are related to the modulation of excitatory synaptic transmission and LTP in the PFC. Pharmaceuticals 16, 288. 10.3390/ph16020288.

45. Pałucha-Poniewiera, A., Podkowa, K., Rafało-Ulińska, A., 2021. The group II mGlu receptor antagonist LY341495 induces a rapid antidepressant-like effect and enhances the effect of ketamine in the chronic unpredictable mild stress model of depression in C57BL/6J mice. Prog. Neuropsychopharmacol. Biol. Psychiatry 109, 110239. 10.1016/j.pnpbp.2020.110239.

46. Pehrson, A.L., Pedersen, C.S., Tølbøl, K.S.., Sanchez, C., 2018. Vortioxetine treatment reverses subchronic PCP treatment-induced cognitive impairments: a potential role for serotonin receptor-mediated regulation of GABA neurotransmission. Front. Pharmacol.9, 162. doi: 10.3389/fphar.2018.00162.

47. Phansalkar, N., More, S., Sabale, A., Joshi, M., 2011. Adaptive local thresholding for detection of nuclei in diversity stained cytology images. In: 2011 International Conference on Communications and Signal Processing. IEEE. 10.1109/iccsp.2011.5739305.

48. Popek, M., Bobula, B., Sowa, J., Hess, G., Frontczak-Baniewicz, M., Albrecht, J., Zielińska, M., 2020. Physiology and morphological correlates of excitatory transmission are preserved in glutamine transporter SN1-depleted mouse frontal cortex. Neuroscience 446, 124–136. 10.1016/j.neuroscience.2020.08.019.

49. Primo, M.J., Fonseca-Rodrigues, D., Almeida, A., Teixeira, P.M., Pinto-Ribeiro, F., 2023. Sucrose preference test: A systematic review of protocols for the assessment of anhedonia in rodents. Eur. Neuropsychopharmacol. 77, 80–92. doi: 10.1016/j.euroneuro.2023.08.496.

50. R Core Team, 2022. R: A language and environment for statistical computing. R Foundation for Statistical Computing, Vienna, Austria. https://www.R-project.org/.

51. Ramsteijn, A.S., Van de Wijer, L., Rando, J., van Luijk, J., Homberg, J.R., Olivier, J.D.A., 2020. Perinatal selective serotonin reuptake inhibitor exposure and behavioral outcomes: A systematic review and meta-analyses of animal studies. Neurosci. Biobehav. Rev. 114, 53–69. 10.1016/j.neubiorev.2020.04.010.

52. Ruggiero, R.N., Rossignoli, M.T., Marques, D.B., de Sousa, B.M., Romcy-Pereira, R.N., Lopes-Aguiar, C., Leite, J.P., 2021. Neuromodulation of Hippocampal-Prefrontal Cortical Synaptic Plasticity and Functional Connectivity: Implications for Neuropsychiatric Disorders. Front. Cell. Neurosci. 15, 732360. 10.3389/fncel.2021.732360.

53. Sanchez, C., Asin, K.E., Artigas, F., 2015. Vortioxetine, a novel antidepressant with multimodal activity: review of preclinical and clinical data. Pharmacol. Ther. 145, 43–57. 10.1016/j.pharmthera.2014.07.001.

54. Sargin, D., Jeoung, H.-S., Goodfellow, N.M., Lambe, E.K., 2019. Serotonin regulation of the prefrontal cortex: cognitive relevance and the impact of developmental perturbation. A.C.S. Chem. Neurosci, 10, 3078−3093. 10.1021/acschemneuro.9b00073.

55. Schendel, D.E., 2017. Prenatal antidepressant use and risk of autism. B.M.J. 358, j3388. 10.1136/bmj.j3388.

56. Schindelin, J., Arganda-Carreras, I., Frise, E., Kaynig, V., Longair, M., et al., 2012. Fiji: an open-source platform for biological-image analysis. Nat. Methods 9, 676–682. 10.1038/nmeth.2019.

57. Schneider, C.A., Rasband, W.S., Eliceiri, K.W., 2012. NIH Image to ImageJ: 25 years of image analysis. Nat. Methods 9, 671–675. 10.1038/nmeth.2089.

58. Silva, C.M., Gonçalves, L., Manhaes-de-Castro, R., Nogueira, M.I., 2010. Postnatal fluoxetine treatment affects the development of serotonergic neurons in rats. Neurosci. Lett. 483, 179–183. 10.1016/j.neulet.2010.08.003.

59. Soiza-Reilly, M., Meye, F.J., Olusakin, J., Telley, L., Petit, E., Chen, X., Mameli, M., Jabaudon, D., Sze, J.-Y., Gaspar, P., 2019. SSRIs target prefrontal to raphe circuits during development modulating synaptic connectivity and emotional behavior. Mol. Psychiatry 24, 726–745. 10.1038/s41380-018-0260-9.

60. Sowa, J., Kusek, M., Siwiec, M., Sowa, J.E., Bobula, B., Tokarski, K., Hess, G., 2018. The 5- HT7 receptor antagonist SB 269970 ameliorates corticosterone-induced alterations in 5- HT7 receptor-mediated modulation of GABAergic transmission in the rat dorsal raphe nucleus. Psychopharmacology 235, 3381–3390. 10.1007/s00213-018-5045-y.

61. Suarez, E.A., Bateman, B.T., Hernández-Díaz, S., Straub, L., Wisner, K.L., et al., 2022. Association of antidepressant use during pregnancy with risk of neurodevelopmental disorders in children. J.A.M.A. Intern. Med. 182, 1149–1160. 10.1001/jamainternmed.2022.4268.

62. Teissier, A., Soiza-Reilly, M., Gaspar, P., 2017. Refining the Role of 5-HT in postnatal development of brain circuits. Front. Cell. Neurosci. 11, 139. 10.3389/fncel.2017.00139.

63. Thase, M.E., Danchenko, N., Brignone, M., Florea, I., Diamand, F., Jacobsen, P.L., Vieta, E., 2017. Comparative evaluation of vortioxetine as a switch therapy in patients with major depressive disorder. Eur. Neuropsychopharmacol. 27, 773–781. 10.1016/j.euroneuro.2017.05.009.

64. Thomas, A., Burant, A., Bui, N., Graham, D., Yuva-Paylor, L.A., Paylor, R., 2009. Marble burying reflects a repetitive and perseverative behavior more than novelty-induced anxiety. Psychopharmacology 204, 361–373. doi: 10.1007/s00213-009-1466-y.

65. Ting, J.T., Lee, B.R., Chong, P., Soler-Llavina, G., Cobbs, C., Koch, C., Zeng, H., Lein, E., 2018. Preparation of acute brain slices using an optimized N-methyl-D-glucamine protective recovery method. J. Vis. Exp. 132, 53825. 10.3791/53825.

66. Unroe, K.A., Maltman, J.L., Shupe, E.A., Clinton, S.M., 2022. Disrupted serotonin system development via early life antidepressant exposure impairs maternal care and increases serotonin receptor expression in adult female offspring. Dev. Psychobiol. 64, e22292. 10.1002/dev.22292.

67. Vitalis, T., Parnavelas, J.G., 2003. The role of serotonin in early cortical development. Dev. Neurosci. 25, 245–256. 10.1159/000072272.

68. Wallace, J., Jackson, R.K., Shotton, T.L., Munjal, I., McQuade, R., Gartside, S.E., 2014. Characterization of electrically evoked field potentials in the medial prefrontal cortex and orbitofrontal cortex of the rat: modulation by monoamines. Eur. Neuropsychopharmacol. 24, 321–332. 10.1016/j.euroneuro.2013.07.005.

69. Yu, W., Yen, Y.-C., Lee, Y.-H., Tan, S., Xiao, Y., Lokman, H., Ting, A.K.T., Ganegala, H., Kwon, T., Ho, W.K., Je, H.S., 2019. Prenatal selective serotonin reuptake inhibitor (SSRI) exposure induces working memory and social recognition deficits by disrupting inhibitory synaptic networks in male mice. Mol. Brain 12, 29. 10.1186/s13041-019-0452-5.

70. Zhong, P., Liu, W., Gu, Z., Yan, Z., 2008a. Serotonin facilitates long-term depression induction in prefrontal cortex via p38 MAPK/Rab5-mediated enhancement of AMPA receptor internalization. J. Physiol., 586, 4465–4479. 10.1113/jphysiol.2008.155143.

71. Zhong, P., Yuen, E.Y., Yan, Z., 2008b. Modulation of neuronal excitability by serotonin-NMDA interactions in prefrontal cortex. Mol. Cell. Neurosci. 38, 290–299. 10.1016/j.mcn.2008.03.003.

